# A Combined Multi-Omics and Supervised Learning Strategy Uncovers an Epithelial Signature of Radiotherapy Response in Colorectal Cancer

**DOI:** 10.1101/2025.04.10.648131

**Authors:** Reuben Kumar, Yujie He, Jiarui Zhou, S:CORT Consortium, Andrew D. Beggs, Deena M.A. Gendoo

**Author notes:** Correspondence: Deena Gendoo Andrew Beggs. Equal Senior Authors (Andrew D. Beggs, Deena M.A. Gendoo). Lead Contact for Submission: Dr. Deena Gendoo.

## Abstract

Colorectal cancer (CRC) is the third most diagnosed cancer globally, accounting for 9.6% of all cancer cases, and the second leading cause of cancer deaths. Radiotherapy is a common treatment, but can demonstrate dangerous side effects and varying patient outcomes that reflects on CRC cancer heterogeneity at genetic, epigenetic, transcriptomics and proteomic levels. Identifying biomarkers capable of effectively predicting CRC patient responses to radiotherapy remains paramount, and an unmet need. We channeled both unsupervised and supervised approaches to assess radiotherapy response for 233 patients of the S:CORT Consortium. Splitting the cohort, we first integrated matched RNA, CNA, mutation, and methylation profiles of 117 patients using multi-omics factor analysis (MOFA). We identified a new radiotherapy signature of 101 biomarkers associated with patients who demonstrate a complete response to radiotherapy, and validated the signature using a random forest classifier on the internal validation dataset, and an independent testing cohort. Our signature effectively predicted treatment outcomes, achieving 89% accuracy with strong discriminatory performance (ROC_AUC = 0.85; PR_AUC = 0.71) to differentiate patients with complete response to radiotherapy compared to incomplete responders. Assessing human and murine scRNAseq datasets underscores that the signature is predominantly expressed in CRC epithelial cells, which underpin CRC heterogeneity and cellular diversity. Our identified signature enables pre-treatment identification of CRC patients that are unlikely to achieve a complete response to radiotherapy, thereby sparing these patients from unnecessary radiation exposure and off-target damage effects.

## BACKGROUND

Colorectal cancer (CRC) encompasses both rectal cancers and colon cancers and is the third most diagnosed cancer globally as of 2022^1^. Accounting for 9.6% of all cancer cases, it is also the second leading cause of cancer deaths, responsible for 9.3% of cancer-related fatalities^1^. Treatments for CRC typically include anterior resection or trans-anal endoscopic microsurgery (TEMS) for early stage rectal cancers. Approximately 50% of rectal patients have locally advanced cancers, and will receive either neoadjuvant chemotherapy or radiotherapy^2^. However, many of these treatments can have significant side effects such as neuropathy, a common consequence of chemotherapy regimens containing oxaliplatin^3^. Treatments involving pelvic radiation can lead to bowel dysfunction including radiation enteritis, perianal irritation, increased stool frequency and incontinence^4,5^. Radiation therapies also have the potential to cause damage to the testicles and ovaries^6,7^. Additionally, radio and chemotherapy can lead to long-term side effects on sexual health and gonadal function^8^. This is exacerbated by increasing rates of CRC incidence among younger individuals in recent decades^9,10^. Such detriments signify the need for personalized treatment strategies, to minimize side effects and to optimize the treatment response, particularly for CRC patients undergoing radiotherapy treatment.

CRC is a heterogeneous disease, with observed cancer heterogeneity at genetic, epigenetic, transcriptomics and proteomic levels^11–14^. Multi-omics approaches can capture facets of the disease, to comprehensively map CRC tumour biology in response to specific treatment regimens. Despite this promise, robust multi-omic analysis to identify molecular features associated with CRC treatment remain underutilized and limited in scope. Yao et al. (2022)^15^ had identified predictive biomarkers of thrombocytopenia (TCP) susceptibility from XELOX chemotherapy treatment in CRC patients, using multi-omics factor analysis (MOFA) to identify key molecular markers associated with treatment, and both LASSO^16^ and SVM classification models^17^ to assess marker efficacy in predicting chemotherapy treatment response. Similarly, another study utilized MOFA to integrate DNA methylation, transcriptomics, and metabolome data from CRC patients undergoing chemotherapy treatment, and identified key biomarkers and pathways associated with hand-foot syndrome^18^. To date however, there is a clear lack of robust, multi-omics analyses of molecular behaviors in CRC that are associated with radiotherapy treatment. Some of these limitations stem from ongoing challenges in integrating multidimensional data, handling missing values associated with different datasets, and selection of effective multi-omic approaches that can address these issues. Recently, Domingo *et al.* conducted a multi-omic analysis of three rectal cancer cohorts, including the Grampian dataset^19,20^, to identify biomarkers predictive of “Complete” patient response to radiotherapy treatment. Combining transcriptomic and genomics data, their work provided initial insights into the biological basis for radiotherapy treatment response CRC, highlighting that immune response features and TGFp signalling play crucial roles in treatment outcome^19^. However, their analysis was restricted to transcriptomic and genomic data, which failed to capture the potential significance of other data modalities, such as epigenetic factors and methylomic alterations on tumour behavior and treatment response^21,22^.

In this work, we develop a robust approach and multi-phased analysis pipeline, channeling a combination of sophisticated machine learning and feature selection techniques applied on multi-omics data, to identify key biomarkers associated with a complete response to radiotherapy treatment in CRC patients. We present the first multi-omic analysis of CRC data using MOFA across four data modalities, including microarray, copy number alteration (CNA), mutational and methylation profiles, trained on a subset of the Grampian cohort^20^. MOFA is unsupervised statistical model with demonstrated ability to handle high dimensionality of multi-omics data^23^ and to extract key features corresponding to specific clinical outcomes (or covariates)^15,24,25^. We identified a panel of 101 biomarkers associated with patients’ response to radiotherapy, and enriched in epithelial cells in CRC patients and disease models. We validated our marker panel using machine learning models on an independent testing cohort from Grampian, and demonstrated 89% predictive ability to differentiate complete patient responders to radiotherapy. Our work identifies markers with significant predictive capability to predict radiotherapy treatment response in CRC, and presents new insights into the mechanisms governing radiotherapy outcomes in CRC patients.

## RESULTS

### Overview of the data and data analysis

We conducted a multi-phase assessment of the Grampian cohort, to identify a predictive panel of patient response to radiotherapy, using multi-omic and supervised machine learning, this panel was then validated on an external dataset (**Figure 1** and see methods, Study Design).

**Figure 1.**
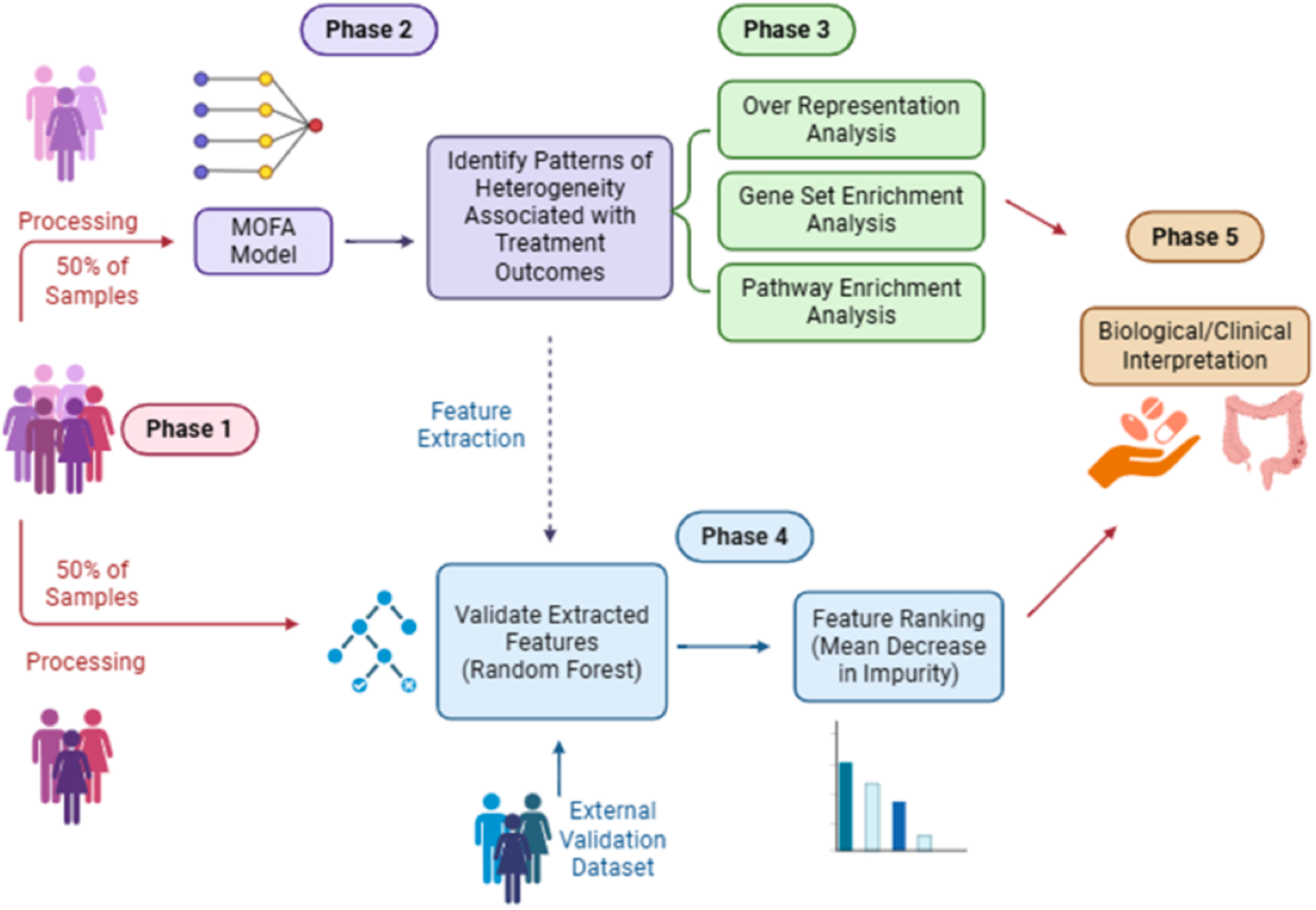
Overview of the data analysis pipeline. The Grampian dataset was split into a training and validation set, and preprocessing of the microarray, CNA, mutational and methylation profiles within each subset was done independently (Phase 1). All four data modalities from the training subset were integrated using MOFA, followed by identifying patterns of variation within the model associated with treatment outcomes (Phase 2). ORA, GSEA and PEA were performed on the identified patterns of variation (Phase 3). Identified features extracted from MOFA were independently verified using random forest models on the validation dataset and separately on an external dataset. Mean decrease in impurity was then used to rank the feature importance within the random forest models (Phase 4). Biological or clinical relevance of the results were further investigated (Phase 5).

Summary plots of the Grampian cohort can be found in **Figure 2** and **Supplementary Figure 1A**, illustrating the distribution of samples against treatment response categories or radiotherapy treatment types, and the number of samples belonging to each data type respectively. Out of 233 patients, only 8 patients were categorized as having a “Minimal” response to radiotherapy treatment, representing 3.43% of the total cohort. The second smallest response category is “Complete” response, containing 37 patients and constitutes 15.87% of the total cohort. When observing the distribution of the treatment types, the “Standard Cap RT” was the most common, spanningl 35 patients or 57.93% of the cohort. The “no concurrent chemo - 50Gy/25#s” treatment type was the least common within this cohort, spanning up 11 patients (4.72%). Across the entire dataset,113 patients were profiled for all 4 data modalities (microarray, mutation, methylation, and CNA profiling). In terms of data modalities, methylation had the lowest representation and microarray had the largest across the cohort, spanning 120 and 225 patients, respectively. We further assessed whether each data modality could capture varying treatment response categories. PCA analysis for each data type (**Supplementary Figure 1B**) indicated that the microarray profiles differentiated samples based on treatment response types, particularly for the “Complete”, “Good Partial” and “Partial” responders.

**Figure 2.**
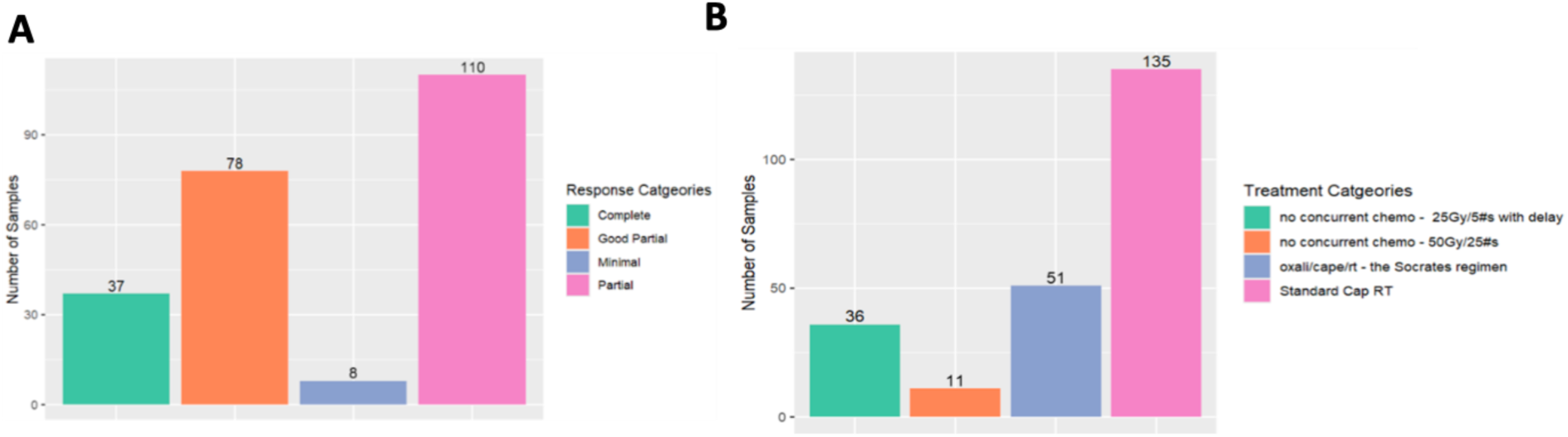
Distribution of treatment response categories and treatment types in the Grampian cohort (a) Barplot of the distributions of samples in different treatment response categories. (b) Barplot of the distribution of samples for radiotherapy treatment types.

### MOFA model selection and development

A total of 117 multi-omic profiles were selected to train the MOFA models, with each model having a differing number of latent factors (LFs) (**Supplementary Figure 2**). The total variance explained captured by the models, for each data modality, demonstrated marked increase as the number of LFs increased, except for the mutation modality which maintained a low variance explained throughout (**Supplementary Figure 2A**). Similarly, the ELBO score increased as the number of LFs increased in the models, and plateaued at 30 LFs (**Supplementary Figure 2B**). For all MOFA models with >20 LFs, each had less than 1 % variance explained in at least one data modality (the least-represented data type). In contrast, the MOFA model with 15 LFs captured 1.65% of the variance explained for the least-represented data modality (**Supplementary Table 1A, 1B**). Accordingly, the MOFA model with 15 latent factors was selected as the most optimal model, and used for the downstream analysis (**Supplementary Figure 3**). We further assessed the correlation between latent factors within this model and determined that none of the latent factors were strongly correlated with each other, confirming that the LFs were not redundant (**Supplementary Figure 3B, Supplementary Table 1A**). This model captured >35% of the variance explained in both the microarray and methylation datasets (35.54% for microarray and 37.28% for methylation respectively) but only captured 16.7% and 0.04% in both the CNA and mutational datasets (**Supplementary Figure 3C**). As the number of latent factors increased, the proportion of variance explained by each additional factor decreased. Accordingly, we observed that variation in the methylation and microarray data is almost exclusively captured within LF1 and LF2 (**Supplementary Figure 3D**).

### Identification of molecular features correlated with treatment response

We probed our MOFA model to investigate the latent factors correlated with treatment response (**Figure 3A, Supplementary Table 2**). Four of the latent factors (LF) were identified, namely latent factor numbers 2,4,5 and 12 (**Figure 3A**). Variance explained for both LF2 and LF4 was predominantly based on microarray data, whereas variance explained by LF5 and LF12 predominantly reflected CNA and methylation variation respectively (**Supplementary Figure 3D)**. Ranking the four factors, the total variance explained by LF2 was the largest (12.93%), followed by LF4 (7.34%), LF5 (5.86%), and LF12 (2.66%) (**Supplementary Table 1B**). Out of these four factors, factor five had the highest correlation to treatment response, with a pearson’s coefficient of 0.40, while factor four had the lowest correlation with a pearson’s coefficient of 0.21 (**Supplementary Table 2**). Interestingly, factor five also showed some correlation with both the overall survival status and disease-free status covariates. Plotting factor values attributed to each sample in the dataset indicated separation between the treatment response categories, most notably for LF2 and LF5 (**Figure 3B**). Using an ANOVA test validated that LF2 and LF5 showed significant differences between treatment response categories, with p-values of 0.0106 and 0.000187 respectively (**Supplementary Table 3**). Plotting samples in the latent factor space underscores the separation of treatment response categories, that reflect on a continuum of response between “Partial”, “Good partial” and “Complete” responders, particularly for latent factors 2, 4, and 5 (**Figure 3C, Supplementary Figure1**).

**Figure 3.**
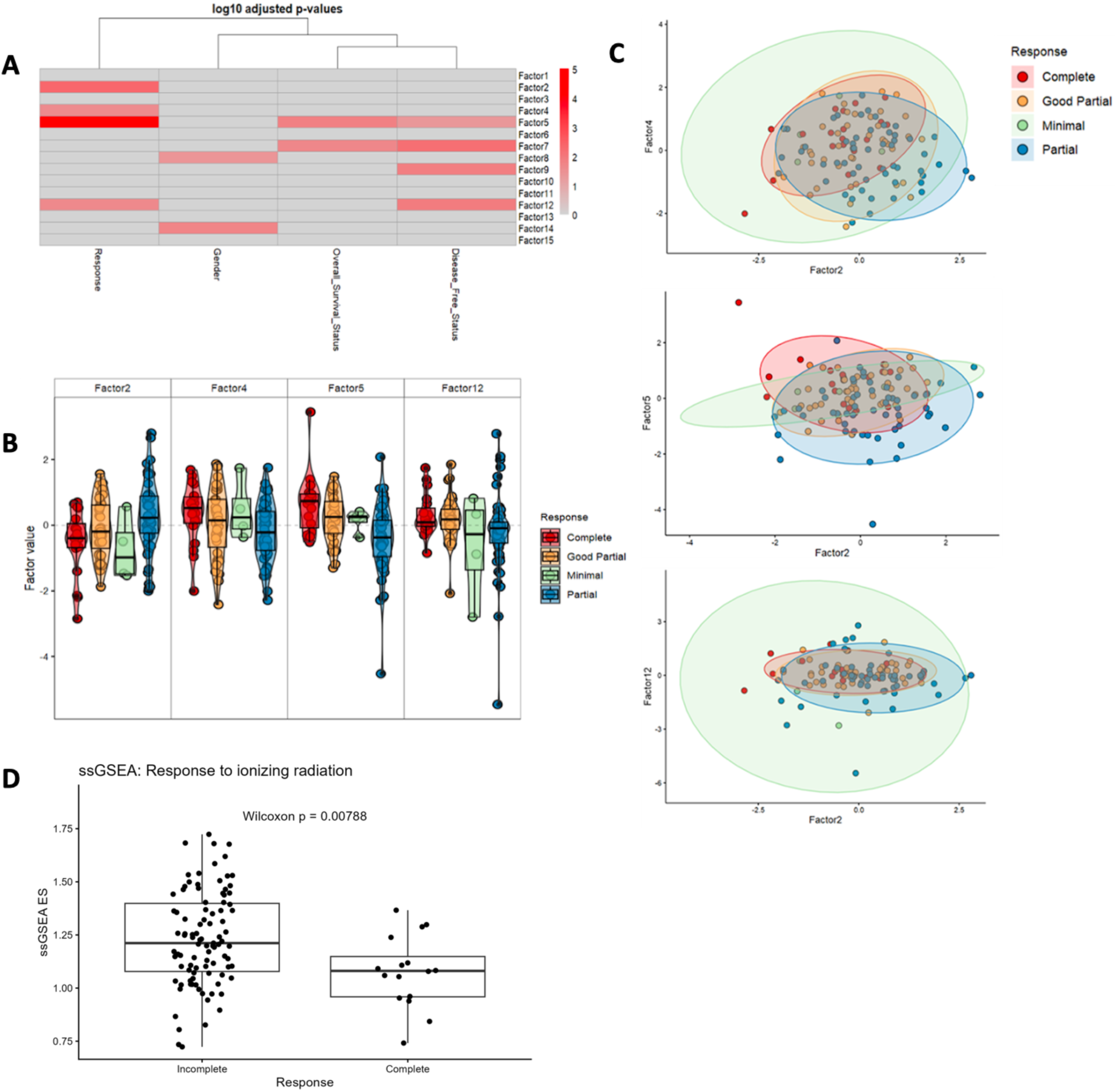
Overview of factor characterization from the MOFA model. (a) Log-adjusted Pearson’s correlation plot between various covariates from patient metadata and the LFs in the MOFA model, the more red the stronger the correlation. (b) Boxplots representing the distribution of different treatment responses in all LFs strongly correlated to treatment response. (c) Samples plotted in 2-dimensional latent factor space across latent factors 2, 4, and 5, colored based on treatment response. (d) ssGSEA analysis using the 5000 genes that were input for the MOFA model, and tested on the validation dataset (111 samples (16 Complete responders, 95 Incomplete responders))

### Biological Pathway Validation of Clinical Phenotypes

As the patient cohorts used in our models includes patients with radiotherapy (RT) and concurrent chemoradiotherapy (CRT), we further probed whether this mixed treatment setting may affect our designated treatment outcomes, specifically complete or incomplete response to radiotherapy. We performed single-sample geneset enrichment analysis (ssGSEA) on our internal validation dataset (n=111 patients) against the KEGG ‘response to ionizing radiation’ pathway, using the 5000 RNA genes that were used as input into our MOFA model. Our analysis demonstrates that our defined complete responders exhibit significantly different pathway activity compared to incomplete responders (**Figure 3D)**, indicating that treatment response is associated with distinct radiation-response biology.

### Extraction of molecular features associated with treatment response defines a signature for radiotherapy response

We exploited our MOFA integrative approach to assess which data modality (or modalities) contributes to predicting radiotherapy response. We assessed the performance of the different data modalities and their features. We first extracted features across a gradient of MOFA loading thresholds. We then evaluated the ability of these features to predict the “Complete” response category using a random forest model on the validation dataset. This approach quickly excluded mutation and copy number profiles, as none of their features exhibited MOFA loadings greater than 0.4, indicating that these data types are not significant and contributed negligibly to the variance explained by the model. Accordingly, we investigated whether a more streamlined RNA and methylation-based model may be present stronger predictive performance. We identified both RNA and methylation features for MOFA loadings between 0.6-0.8 (**Supplementary Figure 4, Supplementary Table4**). Based on the observed PR-AUC and ROC-AUC Curves, a loading threshold of 0.8 suggested optimal performance across the MOFA thresholds tested (**Supplementary Figure 4F**).

We extracted features with loadings >0.8 across the four latest factors correlated with treatment response, across the transcription and methylation profiles (**Supplementary Tables 5A-D, Supplementary Figure 4**). Using this threshold, 101 microarray features and 5 methylation features were extracted. Overall, 78 microarray features pertained to LF2, 23 pertained to LF4, and 5 methylation features pertained to LF12. No features with MOFA loadings >0.8 were observed within LF5. We subsequently questioned the performance of RNA and methylation probes independently, to ascertain which data type was most predictive (**Supplementary Figure 5**). Using methylation probes only demonstrated low discriminatory power when used in a random forest model (ROC AUC 0.54, PR AUC 0.38, **Supplementary Figure 5B**) and were excluded from further analyses. Collectively, these investigations underscore transcriptomics as the only effective data source for predicting radiotherapy response, across the four data types.

To ascertain our feature selection from transcriptomics data, we further queried the discriminatory power of using transcriptomic features alone in the random forest model, across different MOFA thresholds (**Supplementary Figure 5C**). Increasing the threshold from 0.6 to 0.8 led to an improvement in mean PR-AUC from 0.58 (RF0.6) to 0.71 (RF0.8), while the mean ROC-AUC increased from 0.80 to 0.85 over the same range. Further tightening of the threshold to 0.9 or 1.0 did not confer additional gains in ROC AUC (remaining at 0.85 and then decreasing to 0.82) and was accompanied by a reduction in PR AUC to 0.70 and 0.64, respectively. Among all candidate models, RF0.8 therefore exhibited the best overall performance, simultaneously achieving the highest PR AUC and the maximal ROC AUC. On this basis, a MOFA loading threshold of 0.8 was selected as the optimal cut-off, and genes with an absolute loading greater than or equal to 0.8 were defined as the RNA signature.

### Robustness of the Radiotherapy Response Signature

Our radiotherapy response signature consists of 101 genes that were carried forward for subsequent statistical validation and downstream analyses (**Figure 4, Supplementary Figure 6, Supplementary Table 6**). Internal validation of this signature demonstrated a ROC-AUC of 0.85. To account for class imbalance between complete and incomplete responders, we also examined precision-recall (PR) performance, which yielded an average AUC of 0.71 across all folds (**Figure 4A, 4B**). Additionally, the model produced an accuracy score of 0.89 and an F1 score of 0.70 (**Supplementary Table 7A**).

**Figure 4.**
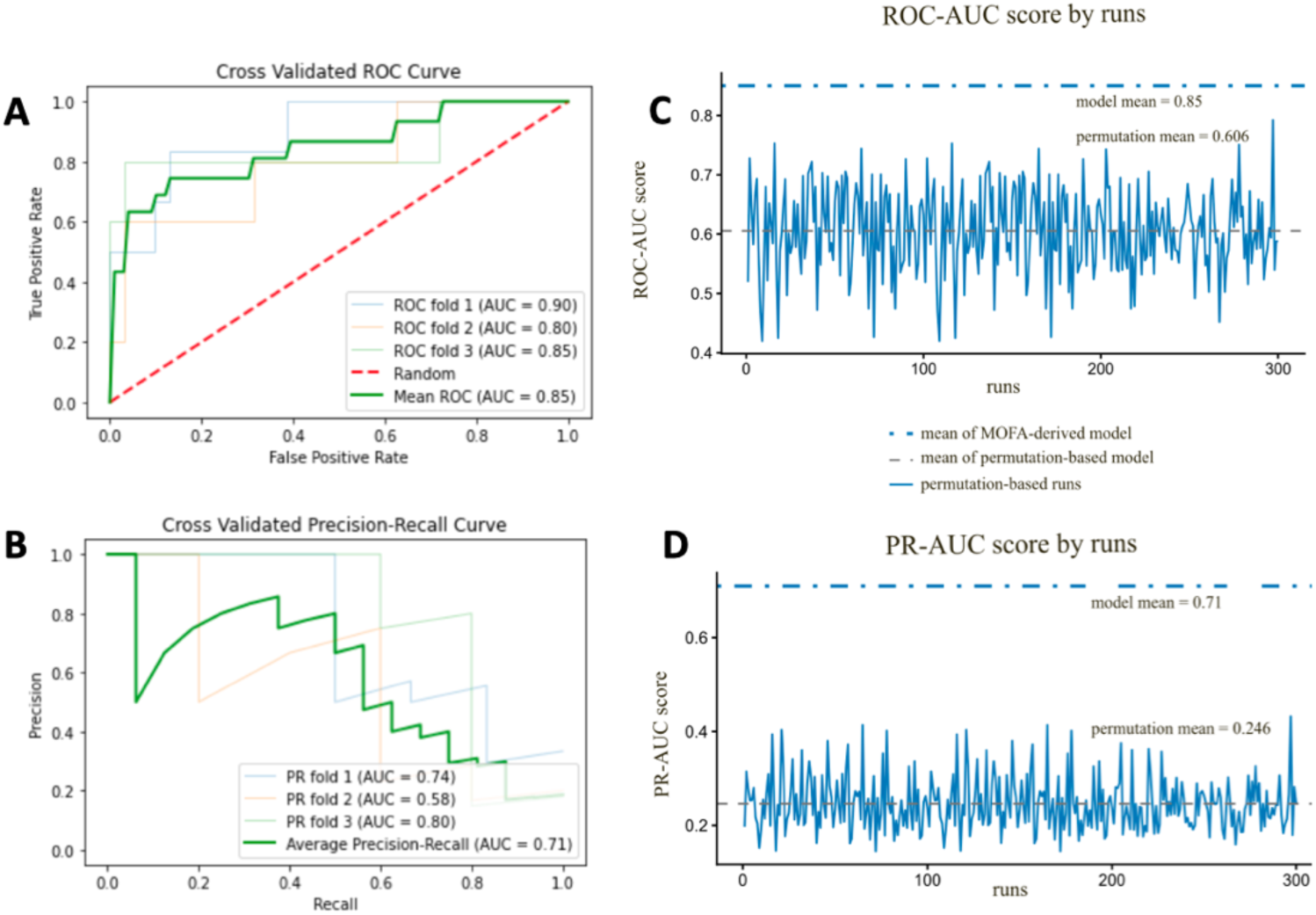
Performance metrics from the random forest model, tested using our 101 microarray features on the validation dataset, with a MOFA loading threshold of 0.8 (a) The receiver operating characteristic curve (ROC curve) for each k-fold, along with an average over all folds. (b) Precision-recall curve (PR curve) for each k-fold, along with an average over all folds. (c) ROC-AUC scores from 300 permutation runs using 101 randomly selected genes under a 3-fold cross-validation scheme. The dashed grey line shows the permutation mean, and the dotted blue line marks the MOFA-derived model mean (ROC-AUC=0.85). (d) PR-AUC scores from the same permutations. The dashed grey line shows the permutation mean, and the dotted blue line marks the MOFA-derived model (PR-AUC=0.71). None of the permutation models exceeded the true model, supporting the robustness of the 101-gene signature.

The robustness of the gene signature was further validated through a 300-run permutation analysis, in which 101 genes were randomly sampled from RNA profiles in the MOFA model in each run, and the performance of the Random Forest classifier was evaluated using a 3-fold cross-validation scheme. The resulting distributions of ROC-AUC and PR-AUC scores showed a marked reduction in predictive performance of random signatures compared with our signature (MOFA-derived model) (**Figure 4C, 4D**). The mean ROC-AUC across all permutation runs was 0.606, whilst the mean PR-AUC reached only 0.246, and none of the randomly generated models exceeded the performance of the true feature-based model when considered across both metrics. These results demonstrate that the predictive capability of the 101-gene panel cannot be attributed to random feature selection, but rather reflect a consistent molecular signature associated with radiotherapy response in colorectal cancer.

As an external validation step, we tested the predictive performance of our signature on RNAseq profiles of an independent, external scRNAseq cohort, for patients of a mixed treatment setting (**Figure 5, Supplementary Table8**). Using these genes as predictors in a random forest classifier evaluated by three-fold cross-validation, the signature retains substantial predictive power in this external dataset (mean ROC-AUC 0.76, mean PR-AUC 0.9, **Figure 5C, 5D**). These findings support that our signature is not susceptible to cohort-specific heterogeneity, but is rather stable and generalizable across datasets.

**Figure 5.**
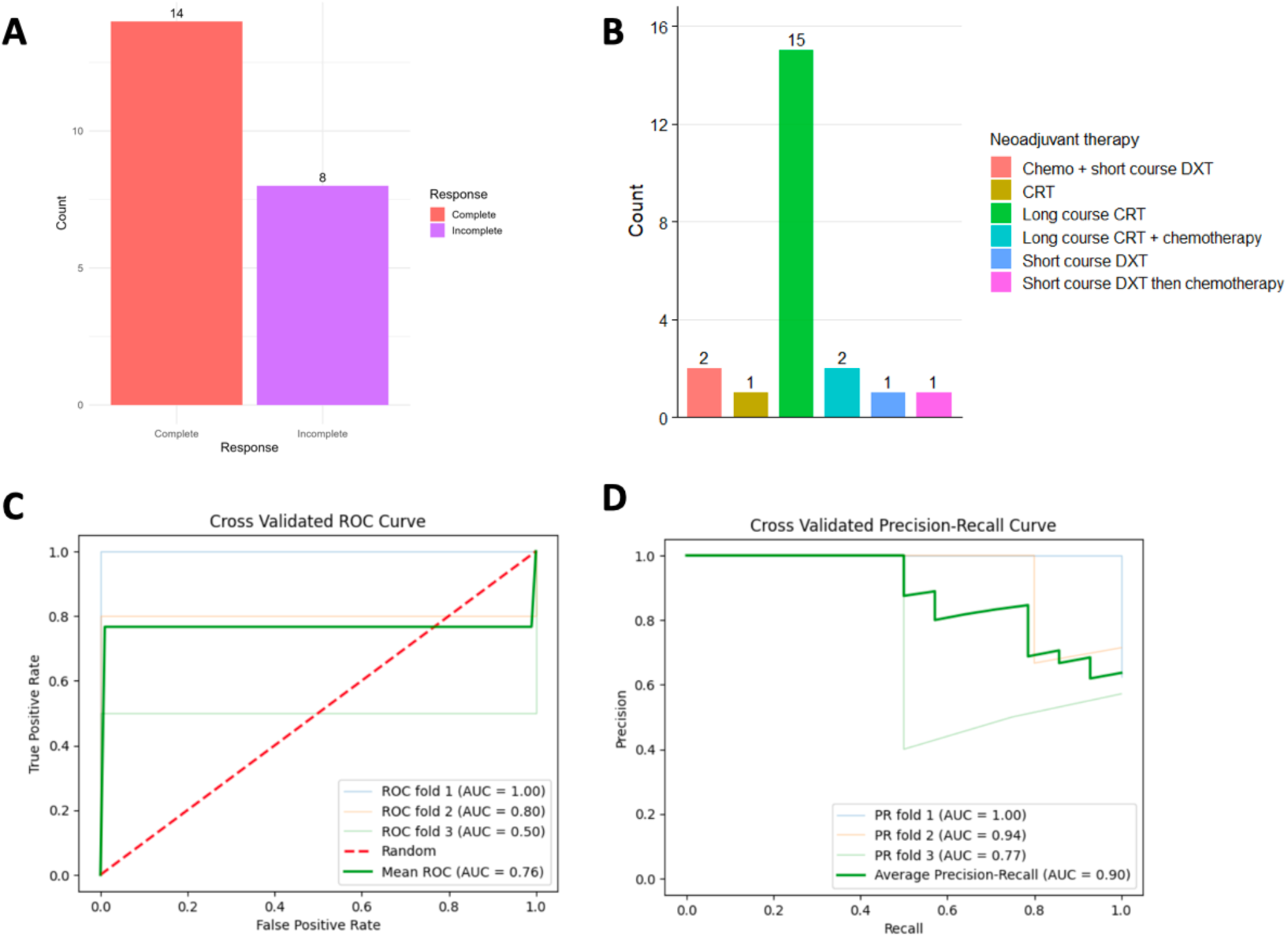
Distribution of treatment response categories and treatment types on the external validation dataset (n=22 samples) (a) Barplot of the distribution of samples for radiotherapy treatment types. (b) Barplot of the distributions of samples in different treatment response categories. (c) External validation of the MOFA-RNA signature in the external dataset. Cross-validated ROC curves of a Random Forest classifier trained on genes selected with a MOFA loading threshold of 0.8, yielding a mean ROC AUC of 0.76. (d) Cross-validated precision-recall curves for the same model, yielding a mean PR AUC of 0.90, indicating that the 0.8-threshold RNA signature retains substantial predictive power for discriminating complete and incomplete responders in the SCORT dataset.

### The signature for radiotherapy response is predominantly expressed in epithelial cells

We further tested our signature for radiotherapy response against several single-cell RNA sequencing datasets, to identify the expression levels of those signature genes across different cell types (**Figure 6, Supplementary Figure 7**). We assessed all 101 genes of the signature and identified that the signature is predominantly expressed in epithelial cells (**Figure 6A**). This observation was corroborated in the analysis of single cells from inducible genetic CRC mouse models, which had been assessed to spatially map cell types and epithelial expression programs to tissue locations in human tumours (**Figure 6B**). We further identified the top 15 contributing features (genes) from our random forest model (**Supplementary Figure 7B**), and ranked the remainder of genes in our signature by mean decrease in impurity (**Supplementary Table 7**). Our analysis demonstrates that the top-performing genes are predominantly expressed in epithelial cells compared to other cells, both in human cancers and mouse models (**Supplementary Figure 7B, 7C**).

**Figure 6.**
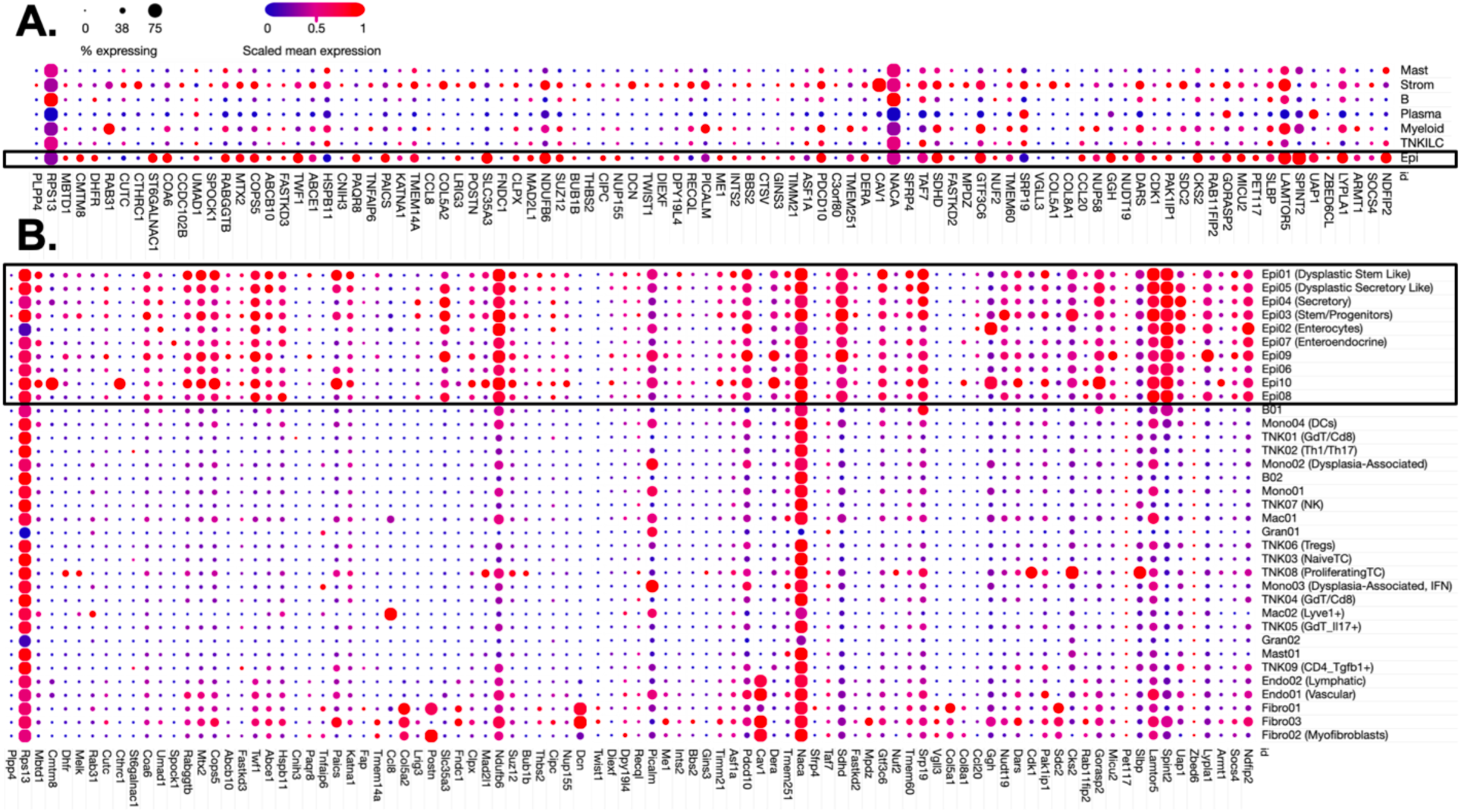
Dotplot representation of the expression levels for 101 genes pertaining to the radiotherapy response signature, assessed across different cell populations from the (a) human colon cancer atlas, and from (b) inducible CRC mouse models. Rows represent cell populations and columns represent individual genes. Circle sizes reflect the percentage of cells expressing a particular gene. Gene expression levels are scaled mean expression, where scaling is relative to each gene’s expression across all cells in a given cell population. Epithelial cell populations are labelled as ‘Epi’ and are highlighted in black boxes.

### Enrichment Analysis

Over-representation analysis (ORA) was performed on the radiotherapy response signature. This analysis revealed enrichment across multiple gene ontology (GO), KEGG, and HALLMARK gene sets. We identified enrichment of several processes related to membrane localization, the cell cycle, mitochondrial components and extracellular matrix-laminin interactions (GO:BP, GO:CC, and KEGG MEDICUS gene sets, **Supplementary Figure 8**). Assessment of HALLMARK gene sets further identified significant enrichment only for E2F targets, which are critical regulators of cell cycle and DNA damage response in CRC, and multicancer invasiveness (MSigDB C2, **Supplementary Figure 8E**). Finally, we conducted a pre-ranked GSEA analysis on GO:CC. We generated a pre-ranked gene list using MOFA model (based on MOFA loadings threshold of 0.8), and extracted MOFA loading weights for the 101 genes of our signature from LF2 and LF4. Pre-ranked GSEA analysis identified significant enrichment for Cell Motility (padj<0.05) (**Supplementary Table9)**.

Gene set analysis (GSEA) and Pathway enrichment analysis (PEA) were also conducted on the top 100 genes extracted from the RNA modality, for each of the 4 latent factors correlated with treatment response of the MOFA model (**Supplementary Figure 9, Supplementary Figure 10, Supplementary Table 10, Supplementary Table 11**). GSEA was performed separately on the positively and negatively loaded transcriptomic features from each of the latent factors. This highlighted positive enrichment of MYC targets (LF 2 and 4, **Supplementary Figure 9a, 9c**) as well as E2F targets (LF2, **Supplementary Figure 9a**) and the G2M checkpoint (LF2, **Supplementary Figure 9a**). Negative enrichment was readily observed for pathways pertaining to PI3K_AKT_MTOR_SIGNALLING, EPITHELIAL_MESENCHYMAL_TRANSITION and TNFA_SIGNALLING_VIA_NFKB signalling (LF, **Supplementary Figure 9b**), as well as hypoxia, TNFA_SIGNALLING_VIA_NFKB and CHOLESTEROL_HOMEOSTASIS (LF5, **Supplementary Figure 9e**).

## DISCUSSION

Channeling a combined unsupervised and supervised approach, our study emphasizes multiple patterns of variation associated with treatment outcomes, and presents a novel and predictive signature associated with radiotherapy treatment response in CRC. This study presents the first comprehensive integration of multi-modal data from CRC patients treated with radiotherapy, providing critical insights into molecular features dictating treatment response outcomes.

Our analysis identified a radiotherapy response signature composed of 101 genes that are predictive of treatment outcomes (**Figure 4-5, Supplementary Table 6, Supplementary Table 7**). We demonstrated that this signature can effectively predict patients who are “Complete” responders to radiotherapy, compared to “Incomplete” responders (limited response or no response), on both internal (**Figure 4**) and external validation cohorts (**Figure 5**). This signature can enable pre-treatment identification of CRC patients that are unlikely to achieve a complete response to radiotherapy, thereby sparing these patients from unnecessary radiation exposure and off-target damage effects. The predictive signature is predominantly associated with epithelial cells in human colon cancers (**Figure 6A**), and similarly, in subtypes of epithelial cells in CRC mouse models (**Figure 6B**). Existing work in CRC and other cancers, by our team and others, underscore the growing use of preclinical models to recapitulate key features of human cancers and in cancer subtyping^26,27^. This agreement across human cancers and murine tissues thus underscores the biological important of the signature in the preclinical and clinical setting, as molecular classification of CRC subtypes has been underpinned by epithelial cellular diversity^28^. Accordingly, our identified epithelial signature presents a useful and predictive panel for early CRC detection of patient response patterns to radiotherapy.

The radiotherapy response signature is associated with biological functions and molecular pathways linked to radiotherapy treatment response and CRC tumorigenesis. A ranked subset of the most predictive features associated with treatment response from the random forest model (**Supplementary Figure 7A, Supplementary Table 7B)** includes several genes associated with CRC tumorigenesis, including *LYPLA1*, *SPINT2*, *LAMPTOR5*, *SLBP*, *PET117*, *MICU2*, *RAB11FIP2*, *CKS2*, and *SDC2* ^29–37^. Assessment of the overall signature also highlights enrichment of the signature for E2F targets, and enrichment of genes related to processes pertaining to membrane localization, the cell cycle, mitochondrial components and extracellular matrix-laminin interactions, and cell mobility and cancer invasiveness (**Supplementary Figure 8**). E2F targets are critical regulators of cell cycle and DNA damage response in CRC, and multicancer invasiveness. E2F transcription factors primarily regulate genes involved in cell proliferation, differentiation and apoptosis through the cell cycle, and are enriched in several cancers^38^. Elevated E2F levels have previously been associated with radiotherapy resistance and poor prognosis^39,40^. Overall, these findings underscore that the biomarker panel includes key determinants that are associated with radiotherapy response and tumour development.

Enrichment analyses across the four response-associated latent factors from MOFA also identified molecular pathways linked to treatment response patterns. GSEA analyses identified upregulation of E2F targets, G2M checkpoint inhibitors, MYC targets, and the PI3K/Akt/mTOR pathway (**Supplementary Figure 9, Supplementary Figure 10**). G2M checkpoints are cell cycle mechanisms that prevent cells with damaged DNA from replicating, and are suggested as a potential target for therapeutic treatments to mediate a more sensitive response to ionising radiation^41,42^. The PI3K/Akt/mTOR pathway is known to promote cell survival, growth and proliferation in response to stimuli^43^ and is commonly dysregulated in cancer^44–46^. In CRC, MYC elevation has been associated with therapeutic resistance via feedback activation of the PI3K/Akt/mTOR pathway^47^. Abnormal activation of this pathway was associated with CRC formation, progression and metastasis^48^. MYC is a proto-oncogene and super-transcription factor that is typically highly regulated but often dysregulated in cancers^49^, and MYC targets were also commonly observed as upregulated across several latent factors. GSEA and PEA analysis also identified negatively regulated pathways associated with treatment response, including Epithelial to Mesenchymal Transition (EMT), a mechanism that facilitates metastases in tumour cells by converting epithelial cells into mesenchymal cells^50^. This transition increases cell mobility and tissue invasion, and has been implicated in chemotherapy and radiotherapy resistance^51^. Changes in TNFA signalling via NFKB were also identified (**Supplementary Figure 9**). NF-KB transcription factors aid in innate and adaptive immune responses, cell death, inflammation and cell proliferation^52^, and their aberrant regulation is associated with cancer development, progression, and resistance to both chemotherapy and radiotherapy^53^. Similarly, identified pathways of hypoxia have previously been strongly associated with radiotherapy resistance^54^. Collectively, these analyses identified recurrent mechanisms that reflect patterns of heterogeneity linked to cancer progression and therapy resistance, and which are also commonly reported in previous studies^19^. This highlights the consistency of these pathways as key plays in CRC development, and their potential as candidate pathways for targeted therapeutic intervention.

To date, this work presents the first multi-modal study to assess and identify a signature associated with CRC radiotherapy outcomes. This approach presented new insight into the data types that are most informative for differentiating patient responses to radiotherapy. MOFA was selected to provide an unbiased and simultaneous comparison of 4 data types (gene expression, methylation, mutation and copy number profiles), to determine the latent biological structure underlying treatment response, and possible axes of variation across the data. In this instance, multi-omics integration served an important role in interpretation and feature prioritization to identify biologically meaningful signal from modality-specific noise. Gene expression (from microarray data) and methylation profiling were the main contributors to the variance captured within the Grampian cohort. This is understandable, given that both data modalities contributed the largest number of features as input to MOFA. The MOFA model captured a continuum of response encapsulated by four latent factors, of which two factors contributed most variation to the model, and both had their variance predominately explained by gene expression profiling. Mutation and copy number profiles contributed a negligible amount of variance to the model, which could reflect high sparsity and low dimensionality within this dataset. However, testing varying MOFA loading thresholds (loadings between 0.4 -1) demonstrated that mutation and copy number features would not have been selected as predictors of treatment response, even at permissive low MOFA loading thresholds (ex: threshold 0.4) (**Supplementary Figure 4**). MOFA followed by supervised learning (random forest) further underscored that across the 4 data types, gene expression was most informative in differentiating patient responses to radiotherapy, compared to methylation profiling (**Supplementary Figure 5, Supplementary Figure 4, Supplementary Figure 1**). A MOFA loading threshold of 0.8 was used to select features that were most predictive of complete response to radiotherapy, including 5 methylation probes and 101 gene expression markers. Notably, methylation profiles are least represented among the Grampian cohort (**Supplementary Figure 1**), so extraction of methylation probes underscores increased methylation heterogeneity within the cohort despite small sample sizes, and reflects on existing findings that DNA methylation is closely linked to CRC^55^. Previous studies have also highlighted the potential of DNA methylation both in enhancing radiation sensitivity in CRC and as predictive biomarkers in neoadjuvant chemoradiotherapy^56,57^. Interestingly however, independent assessment of the gene expression and methylation features demonstrated poor predictive power using methylation probes alone, compared to the 101 gene panel (**Supplementary Figure 5**). Accordingly, our study, as well as Domingo et al.^19^, emphasize transcriptomic biomarkers as stronger predictors of radiotherapy treatment outcomes compared to other data modalities, reinforcing the utility of transcriptomics in clinical decision making.

Our unsupervised and supervised models identified a predictive signature for radiotherapy, using a patient cohort that included individuals who received various treatment regimens, including both radiotherapy (RT) alone and concurrent chemoradiotherapy (CRT). One concern with this approach is that the inclusion of patients treated with both RT and CRT may introduce potential confounding. However, the model was trained to predict treatment response in a real-world clinical setting where patients receive CRT as part of standard-of-care regimens, as chemotherapy is used as a radiosensitizer for RT ^58–60^. As such, the objective of the study was not to isolate a purely radiotherapy-specific mechanistic signature in an RT-controlled system, but to identify predictive biomarkers of response to radiotherapy-based treatment regimens as delivered in practice. To this effect, our signature was also tested on an external cohort which also contains a mixed-treatment setting (**Figure 5**), and the signature still demonstrated discriminatory power. We additionally demonstrated through enrichment analyses (ssGSEA, **Figure 3D**) that despite the composite treatment setting of S:CORT patients, the patient transcriptomic data used for MOFA is strongly linked to radiation response. Our enrichment analysis, both of the signature and the MOFA latent factors, have further identified biological processes and targets associated with RT response. Collectively, these findings support that the model is capturing radiotherapy-specific effects despite a mixed-treatment setting. Accordingly, the signature should be interpreted as reflecting response to radiotherapy-based multimodal treatment, rather than RT in isolation. Future research directions focused on more homogeneous RT-only cohorts are beyond the scope of this study, but would be valuable to further dissect treatment-specific effects.

We observed minimal overlap between our radiotherapy signature against a previously reported panel of complete response predictors by Domingo *et al*, which was also identified using the Grampian cohort^19^. The observed lack of genetic overlap can arise from differences in analytical frameworks, feature selection strategies, and data integration approaches. In this case, a direct head-to-head comparison between both studies is limited by differences in model construction and feature selection. Here, we split the Grampian dataset into a training and validation set, utilizing the cohort for both an unsupervised (MOFA) and supervised (random forest) approach to identify markers associated with treatment. Our approach to integrate multiple data modalities (transcriptome, copy number, methylation, and mutation profiling) further underscored the importance of different data types and their association with treatment response. In contrast, Domingo et al. (2024)^19^ combined the Grampian and Aristotle cohorts as a discovery set, and their analysis was restricted to a transcriptome-based machine-learning workflow. Through a gradient-boosting model, they identified a Radiosensitive Signature (RSS) of 33 genes, capable of predicting response to chemoradiotherapy. Among these 33 genes^19^, only one gene, MICU2, overlaps with our 101-gene signature. This minimal overlap indicates that our approach captures a distinct transcriptional signal associated with radiotherapy response. Despite the lack of overlap, it is worth noting that both signatures highlight complementary and critical biological roles that are crucial for treatment outcomes in CRC. Our GSEA, PEA, and ORA analyses of the MOFA latent factors, as well as the signature itself, highlights processes related to cell cycle regulation and DNA damage response in CRC, and multicancer invasiveness. Domingo *et al*^19^ highlighted that immune response features and TGFp signalling play crucial roles in treatment outcome.

This study has several limitations. One of the challenges relates to an imbalance in the treatment response classes represented within the cohort. Specifically, the ‘Minimal’ response category contained too few samples to be accurately modelled, compared to the ‘Partial’, ‘Good partial’ and “Complete” responders. Despite this shortage for the ‘minimal’ category, the latent factors within MOFA were able to efficiently capture patterns associated with treatment types. Future approaches that binarize treatment categories before MOFA integration may be beneficial to improve classification for the ‘minimal’ response category, although this may impact data granularity. Alternatively, the synthetic minority over-sampling technique (SMOTE) could be considered^61^. Alleviating some of this imbalance by merging patients with minimal, partial, and good partial response also produced a larger number of samples of “Incomplete” responders compared to “Complete” response. Despite this, performance metrics of our supervised machine learning approach indicated that the model could classify the “Complete” response accurately, achieving a robust precision-recall (PR) AUC (**Figure 4**, **Figure 5, Supplementary Table 7**).

Our pipeline has combined both an unsupervised and supervised approach to develop a predictive panel of epithelial biomarkers associated with Complete patient response to radiotherapy treatment. Our analysis also highlighted molecular and biological processes associated with radiotherapy outcomes in CRC. Our predictive panel could be utilized for pre-treatment screening, to assess CRC patients’ response to radiotherapy, enabling more precise and personalized therapeutic strategies. Additionally, this signature can also be utilized towards drug screening of radio-sensitizing agents that potentiate toxicity to radiotherapy, particularly for patients that exhibit “Incomplete” response to treatment.

## METHODS

### Study Design

This project used the Grampian cohort provided by S-CORT^20^, which contains four types of multi- ‘omics profiling for CRC patients: microarray (transcriptomic profiles), methylation profiles, CNA, and mutational profiles. The cohort comprised 233 rectal cancer patients, with corresponding modality profiles generated before radiotherapy treatment. Treatments were one of four types: “standard cap radiotherapy”, “Oxaliplatin/Capecitabine and radiotherapy following the Socrates regimen”, “no concurrent chemo 25Gy/5#s”, and “no concurrent chemo 50Gy/25#s”. All samples were considered in the analysis regardless of treatment type. The patient’s treatment response comprised four categories including: “Minimal” (no reduction in tumour volume or progression), “Partial” (evidence of some response to treatment but substantial tumour remains), “Good Partial” (small foci of residual cancer cells) and “Complete” (complete absence of any microscopic evidence of residual cancer cells).

An overview of the study pipeline is presented in **Figure1**, showing 5 phases of the study. In Phase 1, samples were initially split into two subsets, one for training the unsupervised MOFA model (n=117 samples) and another for validating features extracted from MOFA using a single-omic supervised model (n=116 samples). A stratified 50% split was used to ensure that each subset contained as many samples as possible from each response category. This division was designed to prevent data leakage between the stages. Furthermore, to avoid data leakage, each subset was processed independently. In Phase 2, the four data modalities from the training set were integrated into MOFA to identify patterns of variation associated with treatment outcomes. Gene set enrichment analysis (GSEA) and pathway enrichment analysis (PEA) were performed on identified patterns of variation (Phase 3). Phase 4 validated features from these patterns of variation, by building a random forest model to predict treatment outcomes and rank the importance of each feature on an independent, validation dataset. Biological and clinical interpretation of the results was later performed (Phase 5).

### Microarray Preprocessing

Initially, genes located on sex chromosomes were removed from the microarray dataset. The remaining genes were then log-transformed (log10), and quantile normalized using the R package limma^62^ (version 3.58.1). The data was then subset to only include the top 5000 most varied genes.

### CNA Preprocessing

The CNA dataset was filtered to reduce data sparsity, whereby any features that did not have an observed CNA charge in at least 5% of the samples were removed.

### Mutational Profile Preprocessing

Mutational profiles were filtered to remove sparsity, whereby any mutational events that did not occur in at least 2% of the samples were removed. Given the notable sparsity of this dataset, a less stringent threshold was used compared to the CNA threshold, to preserve mutational events.

### Methylation Preprocessing

Initially, methylation probes on the sex chromosomes were identified using the R package “IlluminaHumanMethylationEPICanno.ilm10b4.hg19” (version 0.6.0) before being removed. The remaining probes were normalized using beta-mixture quantile (BMIQ) normalization from the R package “wateRmelon” (version 2.8.0)^62^ and beta-values were converted to m-values. Finally, data was subset to the top 5000 most varied probes.

### RNA-Seq data analysis

Raw sequencing reads from RNA-Seq experiments were assessed for quality using FastQC software. Sequencing adapter contamination and low-quality bases were then removed from the single end reads using Trim Galore version 0.6.10 ^63^, with a default Phred score cutoff of 20 and automatic trimming of Illumina TruSeq adapter sequence for read one. Processed reads were aligned to the human genome (hg38) with STAR (version 2.7.11b) ^64^ to produce an unsorted BAM file. Genomic alignments were coordinated-sorted using samtools (version 1.21)^65^, and duplicates were identified and marked with Picard MarkDuplicates (version 3.1.1) with default settings. Transcript-level abundances were estimated from the BAM file using Salmon version (1.10.3)^66^ in alignment-based mode.

### Data Visualization

Principal component analysis (PCA) plots were created for each of the pre-processed modality datasets, including all 233 samples, and subsequently labelled with treatment response to gain insight into the overall distribution of the data. To ensure consistency, all datasets were pre-processed in their entirety for this visualization, rather than using individual subsets.

### MOFA Data Integration

Multi-omics factor analysis (MOFA) is an unsupervised statistical model designed to handle the high dimensionality of multi-omics data by identifying common latent factors (LFs) shared across different modality platforms^23^. LFs refer to an underlying pattern representing a dimension of variability across samples. Each feature is given a loading/weight for every LF, similar to a principal component analysis, highlighting the importance of each feature for a given LF. To accomplish this, MOFA uses the following equation:

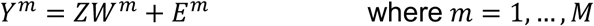

Here, *Y^m^* is the observed data matrix for the modality m, Z is the common LF matrix, *W*^m^ is a weights matrix for each feature in the modality m, *E^m^* is the error matrix and M is the total number of modalities^23^. Importantly, MOFA facilitates the development of multi-omic models even when samples are missing in some datasets, which allows for use of the complete data without the need to impute missing values or remove incomplete samples.

All four modalities within Grampian were integrated using MOFA (version 1.12.1)^23^, with the likelihoods modelled as “bernoulli” for mutation profiles, and “gaussian” for the microarray, CNA, and methylation data. Integration was done using the training subset of samples (n=117 samples). When determining the optimal MOFA model, several models were created and compared. Each model tested a different number of LFs, ranging from 5 to 50 and increasing at intervals of 5. All other parameters remained as default, except “convergence_mode” being set to “slow”, “maxiter” being set to “3000” and “scale_views” being set to “TRUE”, to ensure all modalities were scaled to unit variance. To compare each model’s performance several metrics were considered, namely, the variance explained by each modality within the model, the evidence lower bound (ELBO) that measures how well the model’s approximation of the posterior distribution matches the true distribution of the data, the correlation between each LF, and the minimum percentage of variance explained for at least one modality in each LF. The optimal model was identified as the one with the highest total variance explained and the highest ELBO score while maintaining a low correlation between LFs. However, to ensure the model was not overfitting the data, a threshold was set where all LFs had to explain a minimum of 1% variance in at least one data modality. This set a harsher limit on the number of LFs being considered, to ensure the model would not capture unwanted noise. Following the MOFA model selection stages, a model with 15 LFs was considered for the rest of the analysis.

### Variance Decomposition

The total variance explained per modality within the MOFA model, and the variance captured per LF per modality, were investigated and plotted. This aimed to identify whether the MOFA model was biased towards capturing more variation from certain data type. Understanding how much variance each modality contributed to each LF also determined the relative importance of each modality to specific patterns of heterogeneity within the overall dataset.

### Factor Characterization

Once the optimal MOFA model had been selected, factor characterization was performed using Pearson correlation to identify LFs associated with covariates in the patient metadata. Adjusted p-values from this analysis were plotted and boxplots were generated for LFs that most significantly correlated to the treatment response covariate. These boxplots assessed how each LF captured the distribution of each response category. Samples were plotted in 2-dimensional LF space using these associated LFs and samples were labelled with the treatment response, which can be viewed as a PCA generalized to multiple modalities^25^. This helped ascertain if there are any clustering patterns of the response categories. Subsequently, ANOVA tests were used to identify any significant difference between treatment responses using the factor values given to samples for each LF associated with the response covariate.

### Supervised Learning

Supervised random forest classifier (RFC)^67^ models was produced to classify the samples into “Complete” and “Incomplete” response categories. Samples deemed “Complete” contained only the initial “Complete” response category. Samples classified as “Incomplete” encompassed the “Minimal”, “Partial” and “Good Partial” response categories. Different models were constructed, with feature selection guided by factor loadings across different MOFA loading thresholds (MOFA loading thresholds 0.6, 0.7, 0.8) For a given threshold t, all genes and methylation probes with an absolute loading greater than or equal to t were retained and used as input features to train a Random Forest classifier, to assess if these features could be used as accurate predictors of treatment outcomes. Subsequent testing then assessed the stability and sensitivity of selected features under varying thresholds. For all models, RFC models were constructed and tested on the validation set (n=116 samples), using 70% of these samples to train the RFC and 30% to test the model. Models were constructed using the Python package “Sklearn” (version 1.0.2) using the function “RandomForestClassifier”. Hyperparameter tuning using a grid search cross-validation method was used to find the optimal hyperparameters for the models. To assess model performance, a 3-fold cross-validation approach was employed. Receiver operating characteristic (ROC) and precision-recall (PR) curves, with PR curves included to address class imbalance, were generated along with performance metrics such as the F1 score ^68–70^. These metrics were compared across thresholds in order to identify the loading cut-off that provided the best predictive performance and to define the final threshold to extract the set of MOFA-RNA signature genes.

For final selection of features, models were constructed using features with MOFA loadings > 0.8 from all LFs most significantly correlated with the treatment response. One RFC model was also constructed using transcriptomic features only, and another using methylation features only, to independently assess which data modality was most informative for predicting radiotherapy response.

Following model evaluation, feature importance was assessed using mean decrease in impurity (MDI)^71^ for the transcriptomic model. This allowed features to be ranked based on their relevance in the RFC, suggesting that features with higher ranks are more important in predicting treatment response. Heatmaps were also produced to plot the molecular features with MOFA loadings > 0.8 in each LF, that are most significantly correlated to the treatment response covariate, based on samples belonging to the “Complete” and “Incomplete” response categories. The samples and features were ordered based on hierarchical clustering.

### Permutation Testing

A permutation-based statistical validation analysis was performed to evaluate whether the features extracted from the MOFA model with loadings greater than a predefined threshold represented genuine biological signals, rather than stochastic or noise-driven associations. A total of 300 independent random permutations were conducted. For each iteration, the same number of genes were randomly sampled from the complete pool of genes used in the MOFA model, and were then used as input features to train a Random Forest classifier. Model performance was assessed through a 3-fold cross-validation procedure, and the Receiver Operating Characteristic (ROC) and the Precision-Recall (PR) curves were generated for each permutation. This validation method enabled comparison between the distribution of permutation metrics derived from random feature sets and those obtained from the true MOFA-derived feature set.

### Analysis on single-cell RNA sequencing data

For genes pertaining to the radiotherapy response signature, gene expression levels were assessed in different cell populations using scRNAseq datasets available from the single cell portal (https://singlecell.broadinstitute.org/single_cell). Expression levels were assessed across 371223 cells of the human colon cancer atlas (study SCP1162), which contains several cell types that span B cells, epithelial cells, mast cells, myeloid cells, plasma, stroma, and T, NK, and ILC cell types^72^. Expression levels were also assessed on 915828 cells from inducible genetic CRC mouse models, which contained distinct compositions of cellular subtypes (study SCP1891).

### Over-Representation Analysis

The transcriptomic features extracted from the MOFA model were analyzed using over-representation analysis in R. For this analysis, the 101 features were used as the test gene set, and the 5000 MOFA input genes were used as background. Enrichment analyses were performed with clusterProfiler (version 4.14.6)^73^, using MSigDB Hallmark^74^ and C2 collections^75^ (via msigdbr (version 25.1.1)^76^) and Gene Ontology^77^ (Biological Process, Molecular Function, Cellular Component) categories through enricher() and enrichGO(), respectively. For gene annotation, org.Hs.eg.db (version 3.20.0)^78^ was used. Gene symbols were converted to Entrez IDs using bitr(), and pathway analyses were conducted using KEGG^79^ (enrichKEGG()) and Reactome^80^ (enrichPathway(), ReactomePA (version 1.50.0)^81^. KEGG disease, drug, and network gene sets were retrieved with KEGGREST (version 1.46.0) and tested using enricher() on Entrez IDs. All analyses used the human genome (organism = “hsa”) as the reference, applied Benjamini-Hochberg correction (adjusted *p* < 0.05) and gene set size filters (10-500). Results were visualized using dotplot() from enrichplot(version 1.26.6).

### Pathway Enrichment Analysis

From the LFs most associated with the treatment response covariate, the top 100 genes with the highest MOFA loadings were extracted from the microarray data and used for pathway enrichment analysis (PEA). PEA was performed using the R package “clusterProfiler” (version 4.10.1)^82^ and human gene annotations were provided by the R package “org.Hs.eg.db” (version 3.18.0). Within the function “enrichGO”, three ontologies were considered, namely biological processes (BP), molecular function (MF) and cellular component (CC).

### Gene Set Enrichment Analysis

Gene loadings from the LFs associated with treatment response in the MOFA model were used to perform the gene set enrichment analysis, using the MOFA function “run_enrichment”. Positive and negative enrichment analyses were performed independently and the gene sets used were from “h.all.v2024.1.Hs.symbols.gmt”, taken from the Molecular Signature Database^74^.

### Pre-ranked GSEA and ssGSEA analysis

To generate ranked list of 101 genes for pre-ranked GSEA analysis, we extracted the MOFA loadings for latent factor 2 and factor 4 for the respective genes, with 70 genes from LF2 and 30 genes from LF4. The pre-ranked gene list was compared against the C2-KEGG_LEGACY, C2- REACTOME, and C5-GO:BP genesets from MsigD, as previously described. For single-sample GSEA (ssGSEA), ssGSEA analysis was conducted using the input gene expression matrix for MOFA (5000 genes) and enrichment scores were calculated for each sample using the Gene Ontology (C5) geneset GOBP_RESPONSE_TO_IONIZING_RADIATION (https://www.gsea-msigdb.org/gsea/msigdb/human/geneset/GOBP_RESPONSE_TO_IONIZING_RADIATION.html).

## SUPPLEMENTARY FIGURES

**Supplementary Figure 1. (a)** Venn diagram indicating the availability and overlap of samples across the four data modalities used in this study (RNA (microarray), Mutation, Methylation, and CNA profiles). **(b)** PCA clustering of the Grampian cohort across the different data types, colored based on treatment response.

**Supplementary Figure 2.** MOFA model selection. (a) The total variance explained per data type in a MOFA model, based on increasing numbers of latent factors (b) The Evidence Lower Bound per Factor (ELBO) score of a MOFA model as the number of LFs increases.

**Supplementary Figure 3. Overview of the MOFA model selected for analysis.** (a) Data overview of the MOFA model, illustrating the number of features used for each modality (represented as *D*) for the RNA (microarray), mutation, methylation, and CNA datasets across 117 samples. Grey bars indicate that the sample is missing that type of data. (b) Pearson correlation plot of latent factors in the MOFA model. (c) Total variance explained per modality within the MOFA model. (d) Variance explained per modality, for each factor, within the MOFA model.

**Supplementary Figure 4.** Gene sets and probes were selected from the MOFA RNA/Methylation loadings using a range of thresholds between 0.6-0.8, and used as features for Random Forest modelling in the test dataset. (a) Overview of the number of RNA features extracted for each latent factor, using different MOFA loading thresholds (b) Overview of the number of methylation features extracted for each latent factor, using different MOFA loading thresholds (c) ROC and PR curves showing predictive performance, using 475 genes and 43 CpG probes, based on features extracted using a 0.6 MOFA loading threshold (d) ROC and PR curves showing predictive performance, using 224 genes and 14 CpG probes, based on features extracted using a 0.7 MOFA loading threshold (e) ROC and PR curves showing predictive performance, using 101 genes and 5 CpG probes, based on features extracted using a 0.8 MOFA loading threshold (f) Overall summary of the mean PR AUC and mean ROC AUC across the different MOFA thresholds.

**Supplementary Figure 5.** Predictive performance of RNA and methylation probes independently, using the MOFA threshold 0.8. (a) ROC and PR curves showing predictive performance, using 101 genes (microarray features) (b) ROC and PR curves showing predictive performance, using 5 CpG probes (c) Performance of Random Forest models trained with MOFA-RNA features selected at different loading thresholds between 0.6-1. Models was built using genes whose absolute MOFA loading exceeded the indicated cut-off. Mean area under the ROC curve (ROC AUC, dashed line with triangles) and mean area under the precision-recall curve (PR AUC, solid line with circles) were estimated by cross-validation. The model based on a loading threshold of 0.8 (RF0.8) achieved the highest overall performance, with a mean ROC AUC of 0.85 and a mean PR AUC of 0.71, and was therefore selected for definition of the RNA signature.

**Supplementary Figure 6.** Heatmap of microarray expression values. A heatmap of the 101 microarray features extracted from MOFA are plotted for samples of the validation dataset, with samples stratified based on response (Complete or Incomplete response).

**Supplementary Figure 7:** The radiotherapy response signature is predominantly expressed in epithelial cell populations (a) Mean decrease in impurity plot, showing the ranking of feature importance within the random forest model and the top 15 genes that contribute to the predictive signature (b-c) Dotplot representation of the expression levels for the top 15 genes from the radiotherapy signature. Genes are selected based on ranking feature importance from the random forest model. Expression levels are plotted against different cell populations for (b) human colon cancers and (c) inducible genetic mouse models. Rows represent cell populations and columns represent individual genes. Circle sizes reflect the percentage of cells expressing a particular gene. Gene expression levels are scaled mean expression, where scaling is relative to each gene’s expression across all cells in a given cell population. Cell populations pertaining to Epithelial cells are labelled as ‘Epi’.

**Supplementary Figure 8.** Dotplots for over-representation analyses on MOFA extracted genes. ORA conducted on the 101 genes extracted from the MOFA model based on associations to treatment response. (a) Dotplot assessing enrichment for gene ontology biological processes. (b) Dotplot assessing enrichment for gene ontology cellular components. (c) Dotplot assessing enrichment for gene ontology molecular functions. (d) Dotplot assessing enrichment for MSigDB hallmarks. (e) Dotplot assessing enrichment for MSigDB C2 collections. (f) Dotplot assessing enrichment for KEGG medicus.

**Supplementary Figure 9.**Gene set enrichment analysis (GSEA) of MOFA LFs correlated with treatment response. GSEA was performed separately on the positively and negatively loaded RNA features from the four LFs correlated to treatment response. Gene sets analysed were “Hallmark Gene Sets” from MSigDB. LFs 2,4 and 12 exhibited enrichment (a-c), whereas for the negatively loaded features only LFs 4 and 5 showed enrichment (d-e). The only significantly enriched gene set for LF12 was MYC targeting v1(c).

**Supplementary Figure 10.** Pathway enrichment analysis. Dot plots displaying significantly enriched pathways from the RNA features taken from MOFA LFs correlated to treatment response. Dot size indicates gene counts enriched in a pathway and colour represents adjusted p-values, with red showing higher significance. (a) Biological process ontology. (b) Molecular function ontology. (c) Cellular component ontology.

## SUPPLEMENTARY TABLES

**Supplementary Table 1. (A)** Pearson correlation coefficients between LFs. **(B)** Variance explained by each LF for each omics dataset within the optimal MOFA model.

**Supplementary Table 2. (A)** Pearson correlation coefficients assessing the relationship between each latent factor from the MOFA model, and patient metadata covariates. **(B)** log10 adjusted p-values from the Pearson correlation analysis, correlating patient metadata with factors in the MOFA model.

**Supplementary Table 3**. Results of ANOVA tests evaluating significant differences between treatment response types based on factor values assigned to each sample.

**Supplementary Table 4.** Transformed read counts for the transcriptomic (gene) features and methylation probes identified at different MOFA loading thresholds between 0.6 and 0.8.

**Supplementary Table 5.** MOFA weights/loadings of all features for each data modality, based on MOFA loading threshold 0.8.

**Supplementary Table 6.** All features extracted from the MOFA model with loadings >0.8.

**Supplementary Table 7.** Performance of the Random Forest model. (A) Performance metrics of the random forest model. (B) MDI scores for features in the random forest model.

**Supplementary Table 8.** The External Validation dataset (22 samples). (A) Clinical characteristics of the patient cohort. The “prognostic group” column was used to label patients for complete and incomplete response. (B) Gene count matrix following scRNAseq data processing. (C) List of genes in the radiotherapy signature that were tested in the external dataset

**Supplementary Table 9.** Pre-ranked GSEA Analysis. (A) Ranked gene list based on MOFA loadings (B) Pre-ranked GSEA on the MSigDB C5 gene sets (C) Pre-ranked GSEA on the MSigDB C2 gene sets.

**Supplementary Table 10.** Gene set enrichment analysis (GSEA) using the top 100 features from factors most correlated to treatment response, to identify significantly enriched pathway from the MSigDB Hallmark gene set. GSEA was conducted on positive loadings (A) and negative loadings (B).

**Supplementary Table 11**. Pathway enrichment analysis (PEA) using the top100 features from factors most correlated to treatment response, to identify significantly enriched pathways from Gene Ontology, using GO gene sets for (A) molecular function, (B) biological pathways, and (C) cellular components.

## Author Contributions

Conceived the study: A.D.B, D.M.A.G

Bioinformatics data analysis and interpretation: R.K, Y.H., D.M.A.G

Development of computational methodology: R.K, Y.H., D.M.A.G

Review of Study Design: J.Z

Code Development: R.K, Y.H.

Study Supervision: D.M.A.G

Data Acquisition: A.D.B

Writing, review, and/or revision of the manuscript: R.K, Y.H., A.D.B, D.M.A.G

A.D.B and D.M.A.G are joint co-senior authors on this work.

Approved the manuscript: all authors

## Acknowledgements

The authors would like to thank Dr. Venkata Manem (Laval University and CHU de Quebec) for advice on future directions of this work.

