## Supplementary Figures 1-10 for "A Combined Multi-Omics and Supervised Learning Strategy Uncovers an Epithelial Signature of Radiotherapy Response in Colorectal Cancer"

Reuben Kumar[1], Yujie He[1], Jiarui Zhou[2], S:CORT Consortium, Andrew D. Beggs[1,$,*], Deena M.A. Gendoo[1,3,$,*]

1. Department of Cancer and Genomic Sciences, School of Medical Sciences, College of Medicine and Health, University of Birmingham, B15 2TT, United Kingdom.

2. School of Biosciences and Centre for Environmental Research and Justice (CERJ), University of Birmingham, Birmingham, B15 2TT, United Kingdom

3. Institute for Data and AI, University of Birmingham, Birmingham, United Kingdom

$ Equal Senior Authors

* Correspondence:

Deena Gendoo

Andrew Beggs;

Lead Contact for Submission:

Dr. Deena Gendoo,

#### Supplementary Figure 1

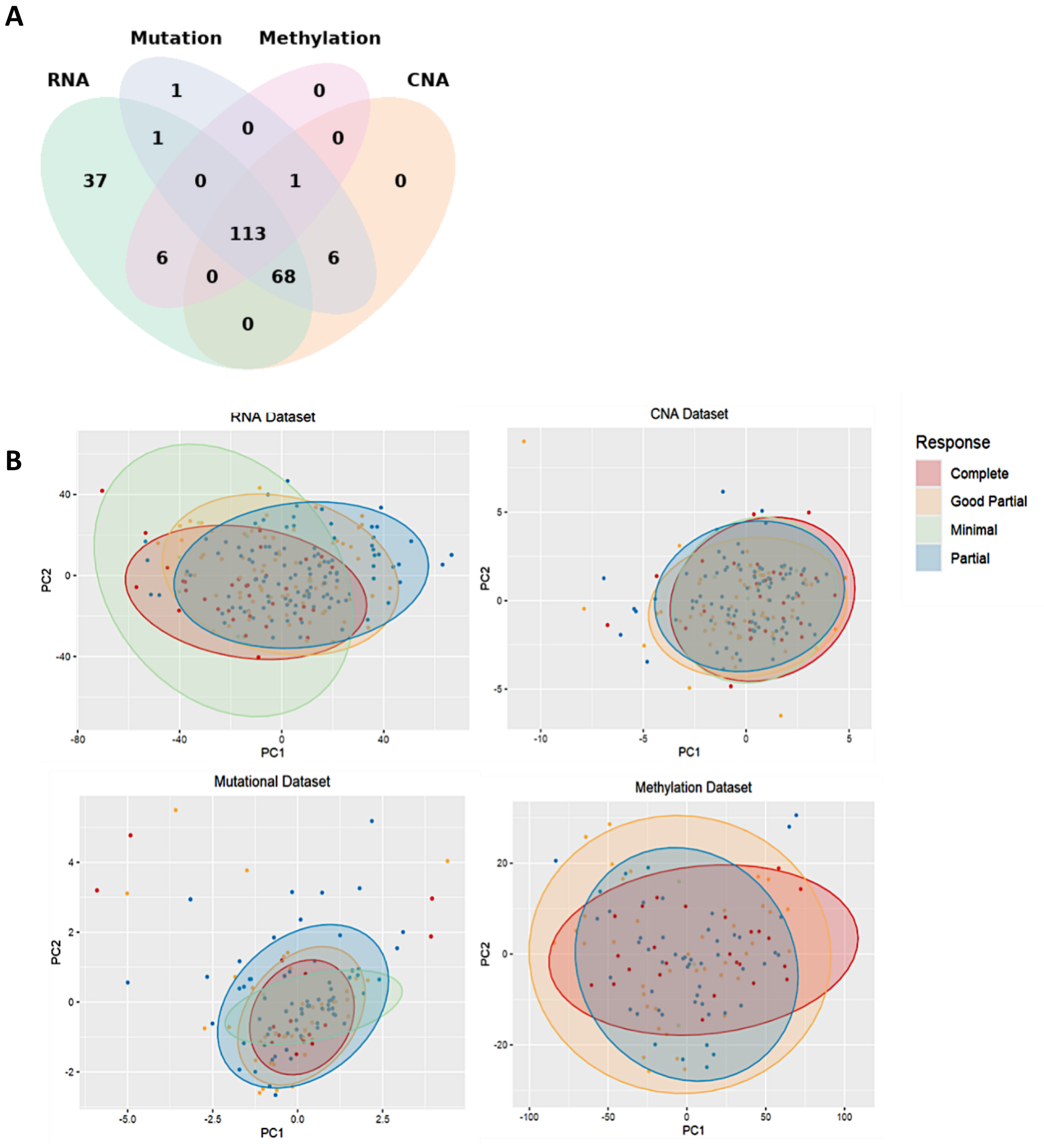


**Supplementary Figure 1**.

(a) Venn diagram indicating the availability and overlap of samples across the four data modalities used in this study (RNA (microarray), Mutation, Methylation, and CNA profiles).

(b) PCA clustering of the Grampian cohort across the different data types, colored based on treatment response.

**A Combined Multi-Omics and Supervised Learning Strategy Uncovers an Epithelial Signature of Radiotherapy Response in Colorectal Cancer**

Reuben Kumar, Yujie He, Jiarui Zhou, S:CORT Consortium, Andrew D. Beggs, Deena M.A. Gendoo

#### Supplementary Figure 2

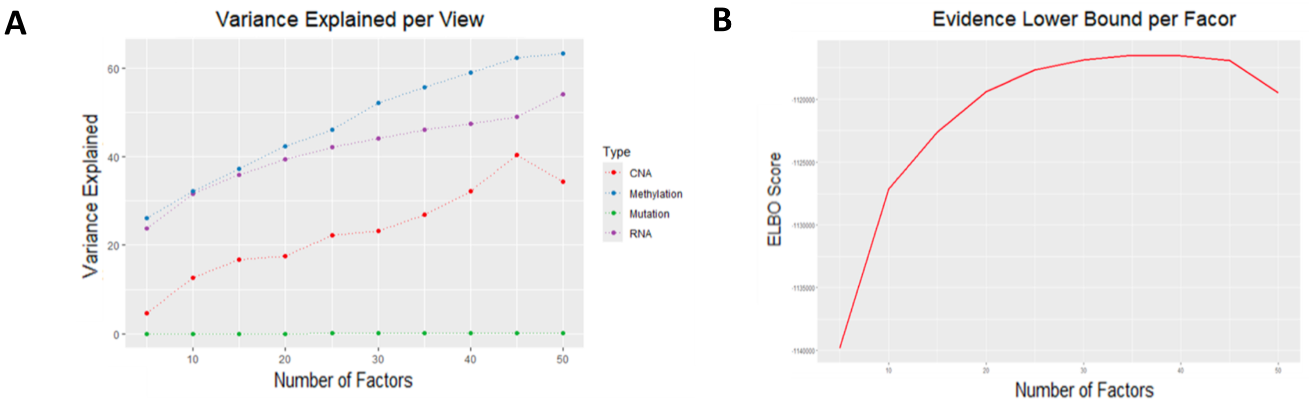


**Supplementary Figure 2.** MOFA model selection.

(a) The total variance explained per data type in a MOFA model, based on increasing numbers of latent factors

(b) The Evidence Lower Bound per Factor (ELBO) score of a MOFA model as the number of LFs increases.

**A Combined Multi-Omics and Supervised Learning Strategy Uncovers an Epithelial Signature of Radiotherapy Response in Colorectal Cancer**

Reuben Kumar, Yujie He, Jiarui Zhou, S:CORT Consortium, Andrew D. Beggs, Deena M.A. Gendoo

#### Supplementary Figure 3

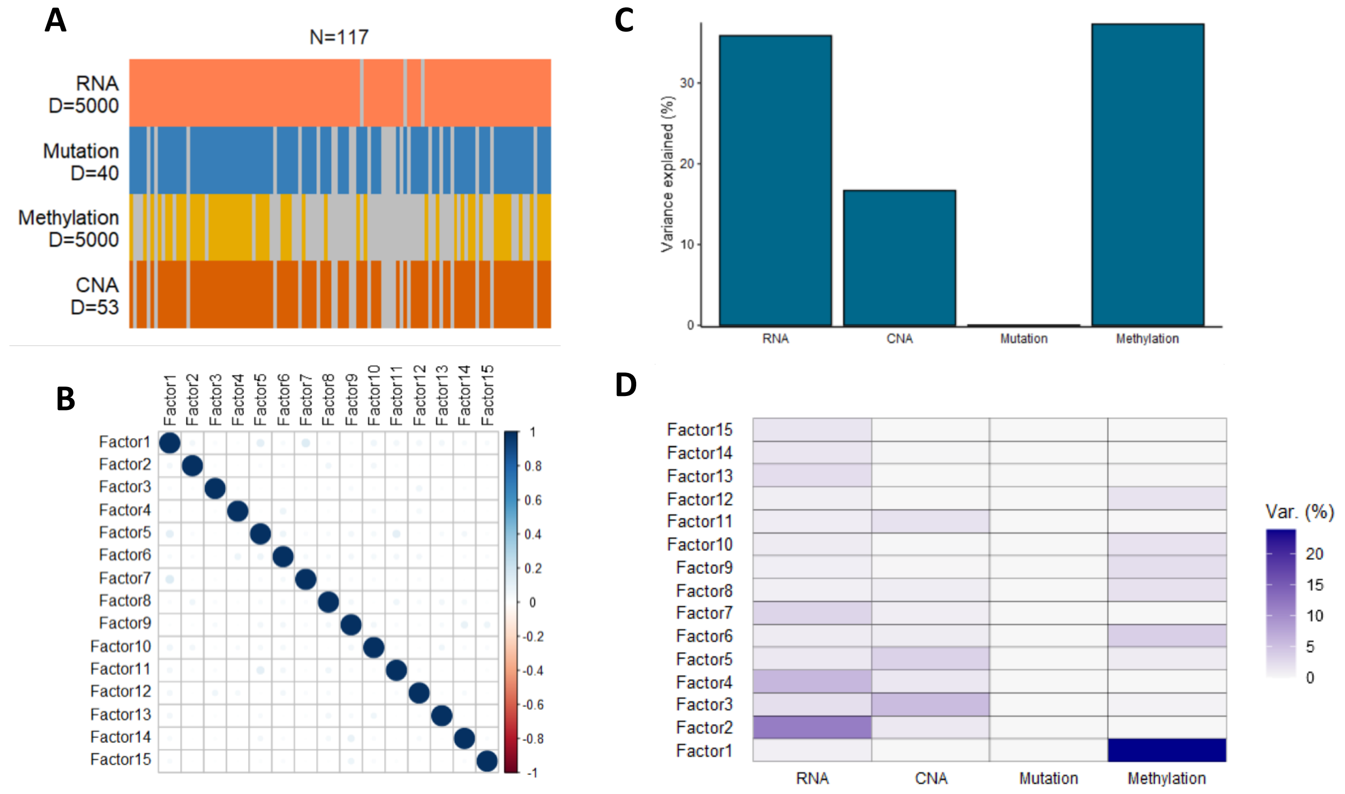


**Supplementary Figure 3.** Overview of the MOFA model selected for analysis

(a) Data overview of the MOFA model, illustrating the number of features used for each modality (represented as *D*) for the RNA (microarray), mutation, methylation, and CNA datasets across 117 samples. Grey bars indicate that the sample is missing that type of data.

(b) Pearson correlation plot of latent factors in the MOFA model.

(c) Total variance explained per modality within the MOFA model.

(d) Variance explained per modality, for each factor, within the MOFA model.

### Supplementary Figure 4

**A** RNA features

| | threshold | | | | | |
|---|---|---|---|---|---|---|
| factor | 0.4 | 0.6 | 0.7 | 0.8 | 0.9 | 1 |
| 2 | 995 | 336 | 162 | 78 | 34 | 11 |
| 4 | 433 | 142 | 62 | 23 | 12 | 5 |
| 5 | 0 | 0 | 0 | 0 | 0 | 0 |
| 12 | 0 | 0 | 0 | 0 | 0 | 0 |

**B** Methylation features

| | threshold | | | | | |
|---|---|---|---|---|---|---|
| factor | 0.4 | 0.6 | 0.7 | 0.8 | 0.9 | 1 |
| 2 | 0 | 0 | 0 | 0 | 0 | 0 |
| 4 | 0 | 0 | 0 | 0 | 0 | 0 |
| 5 | 0 | 0 | 0 | 0 | 0 | 0 |
| 12 | 396 | 43 | 14 | 5 | 0 | 0 |

**C**

MOFA 0.6
475 genes
43 CpG probes

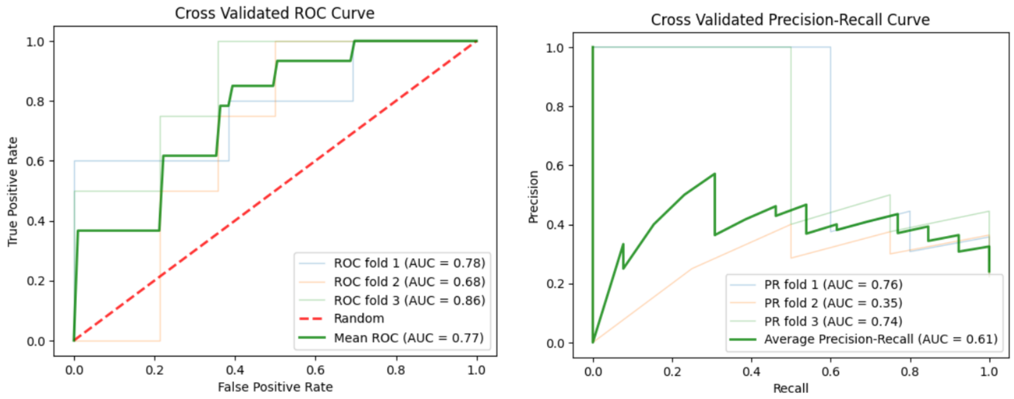


**D**

MOFA 0.7
224 genes
14 CpG probes

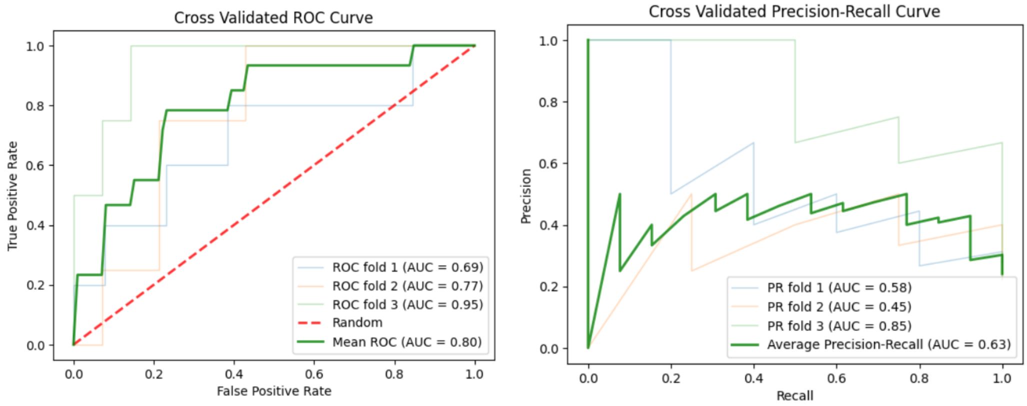


**E**

MOFA 0.8
101 genes
5 CpG probes

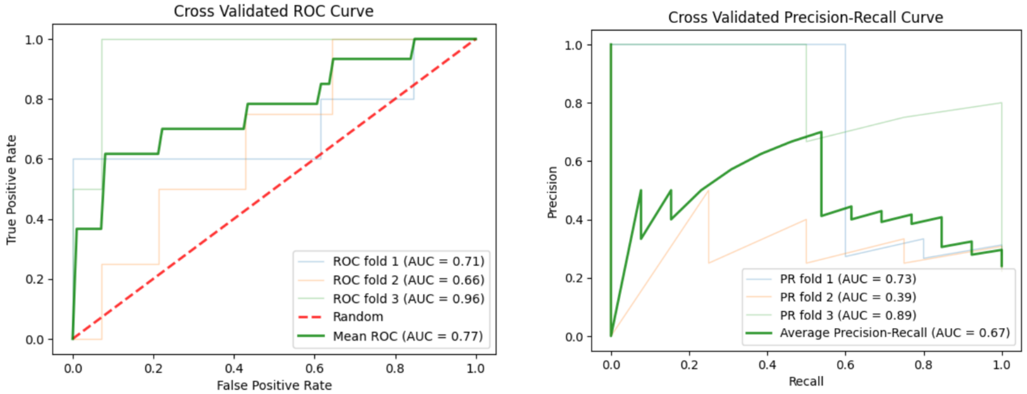


**F**

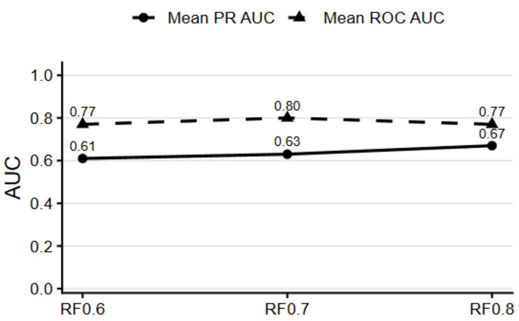


**Supplementary Figure 4.** Gene sets and probes were selected from the MOFA RNA/Methylation loadings using a range of thresholds between 0.6-0.8, and used as features for Random Forest modelling in the test dataset. (a) Overview of the number of RNA features extracted for each latent factor, using different MOFA loading thresholds (b) Overview of the number of methylation features extracted for each latent factor, using different MOFA loading thresholds (c) ROC and PR curves showing predictive performance, using 475 genes and 43 CpG probes, based on features extracted using a 0.6 MOFA loading threshold (d) ROC and PR curves showing predictive performance, using 224 genes and 14 CpG probes, based on features extracted using a 0.7 MOFA loading threshold (e) ROC and PR curves showing predictive performance, using 101 genes and 5 CpG probes, based on features extracted using a 0.8 MOFA loading threshold (f) Overall summary of the mean PR AUC and mean ROC AUC across the different MOFA thresholds.

**A Combined Multi-Omics and Supervised Learning Strategy Uncovers an Epithelial Signature of Radiotherapy Response in Colorectal Cancer**

Reuben Kumar, Yujie He, Jiarui Zhou, S:CORT Consortium, Andrew D. Beggs, Deena M.A. Gendoo

#### Supplementary Figure 5

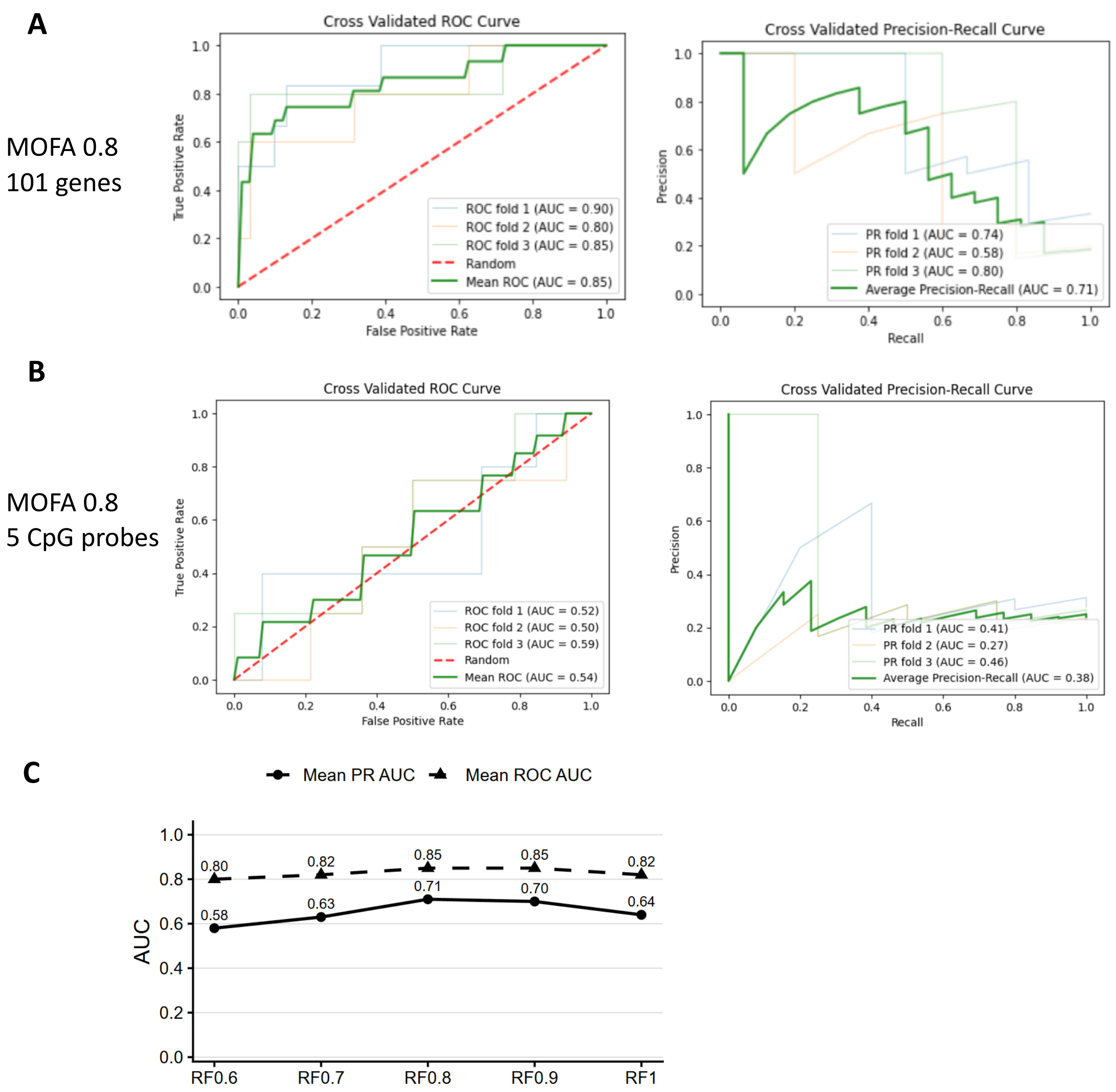


**Supplementary Figure 5. Predictive performance of RNA and methylation probes independently, using the MOFA threshold 0.8.** (a) ROC and PR curves showing predictive performance, using 101 genes (microarray features) (b) ROC and PR curves showing predictive performance, using 5 CpG probes (c) Performance of Random Forest models trained with MOFA-RNA features selected at different loading thresholds between 0.6-1. Models was built using genes whose absolute MOFA loading exceeded the indicated cut-off. Mean area under the ROC curve (ROC AUC, dashed line with triangles) and mean area under the precision–recall curve (PR AUC, solid line with circles) were estimated by cross-validation. The model based on a loading threshold of 0.8 (RF0.8) achieved the highest overall performance, with a mean ROC AUC of 0.85 and a mean PR AUC of 0.71, and was therefore selected for definition of the RNA signature.

**A Combined Multi-Omics and Supervised Learning Strategy Uncovers an Epithelial Signature of Radiotherapy Response in Colorectal Cancer**

Reuben Kumar, Yujie He, Jiarui Zhou, S:CORT Consortium, Andrew D. Beggs, Deena M.A. Gendoo

#### Supplementary Figure 6

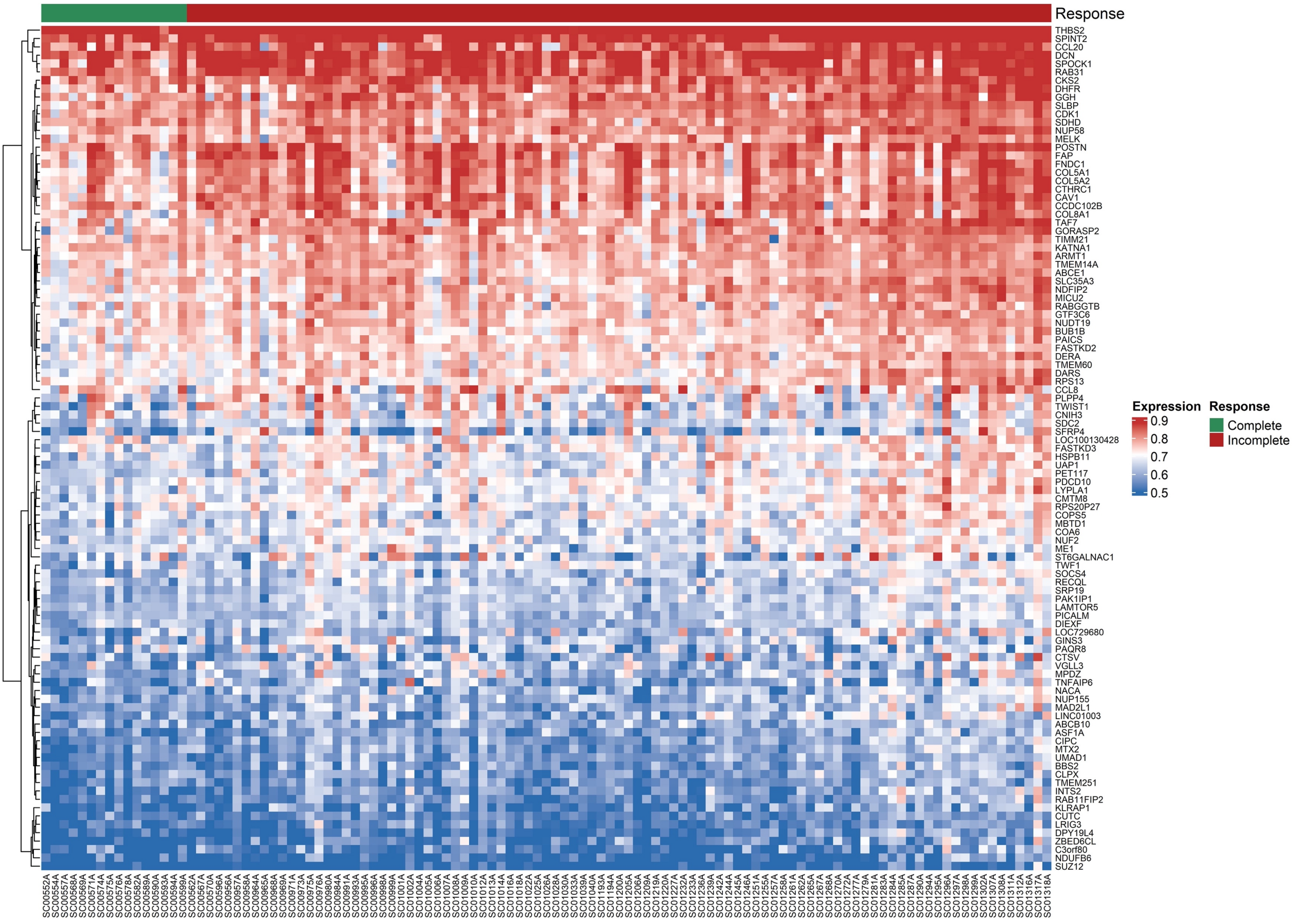


**Supplementary Figure 6**. Heatmap of microarray expression values. A heatmap of the 101 microarray features extracted from MOFA are plotted for samples of the validation dataset, with samples stratified based on response (Complete or Incomplete response).

#### Supplementary Figure 7

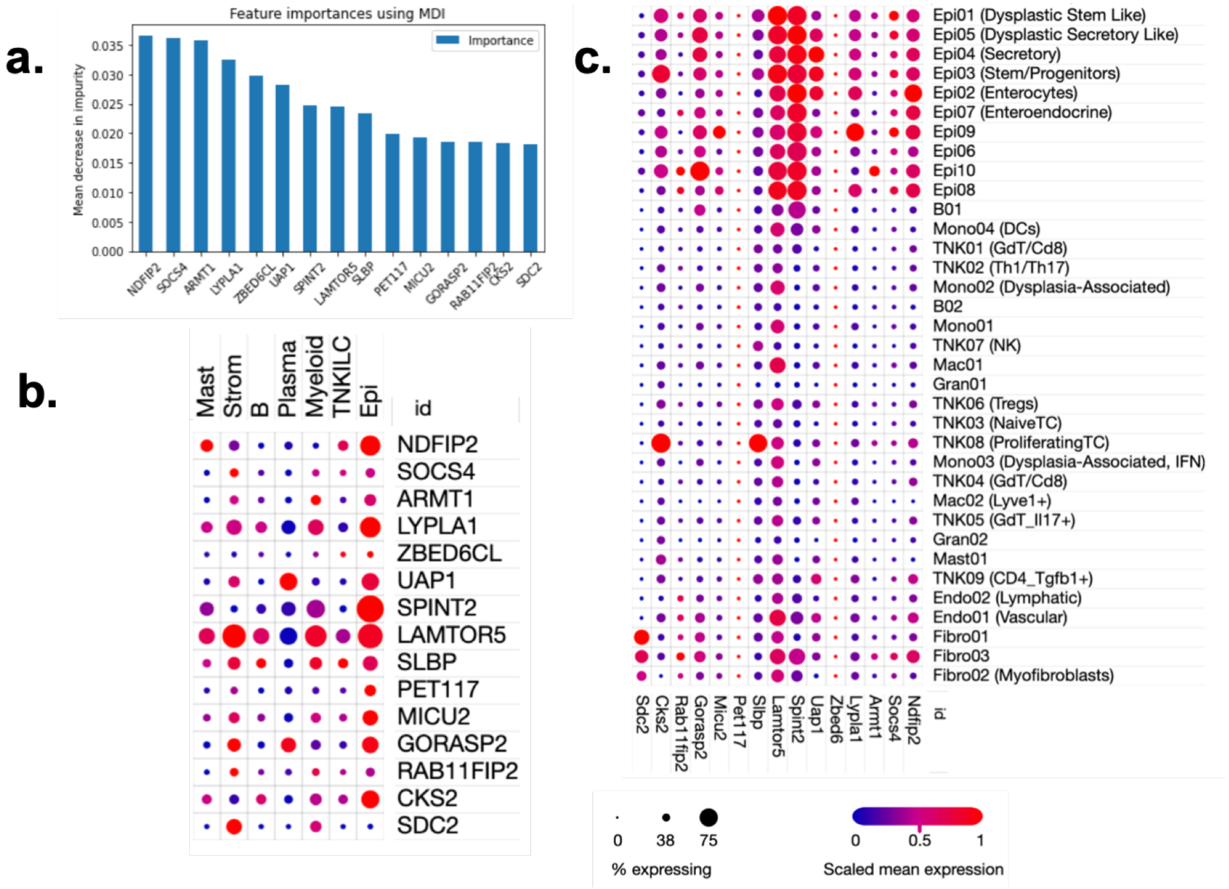


**Supplementary Figure 7: The radiotherapy response signature is predominantly expressed in epithelial cell populations** (a) Mean decrease in impurity plot, showing the ranking of feature importance within the random forest model and the top 15 genes that contribute to the predictive signature (b-c) Dotplot representation of the expression levels for the top 15 genes from the radiotherapy signature. Genes are selected based on ranking feature importance from the random forest model. Expression levels are plotted against different cell populations for (b) human colon cancers and (c) inducible genetic mouse models. Rows represent cell populations and columns represent individual genes. Circle sizes reflect the percentage of cells expressing a particular gene. Gene expression levels are scaled mean expression, where scaling is relative to each gene's expression across all cells in a given cell population. Cell populations pertaining to Epithelial cells are labelled as 'Epi'.

**A Combined Multi-Omics and Supervised Learning Strategy Uncovers an Epithelial Signature of Radiotherapy Response in Colorectal Cancer**

Reuben Kumar, Yujie He, Jiarui Zhou, S:CORT Consortium, Andrew D. Beggs, Deena M.A. Gendoo

#### Supplementary Figure 8

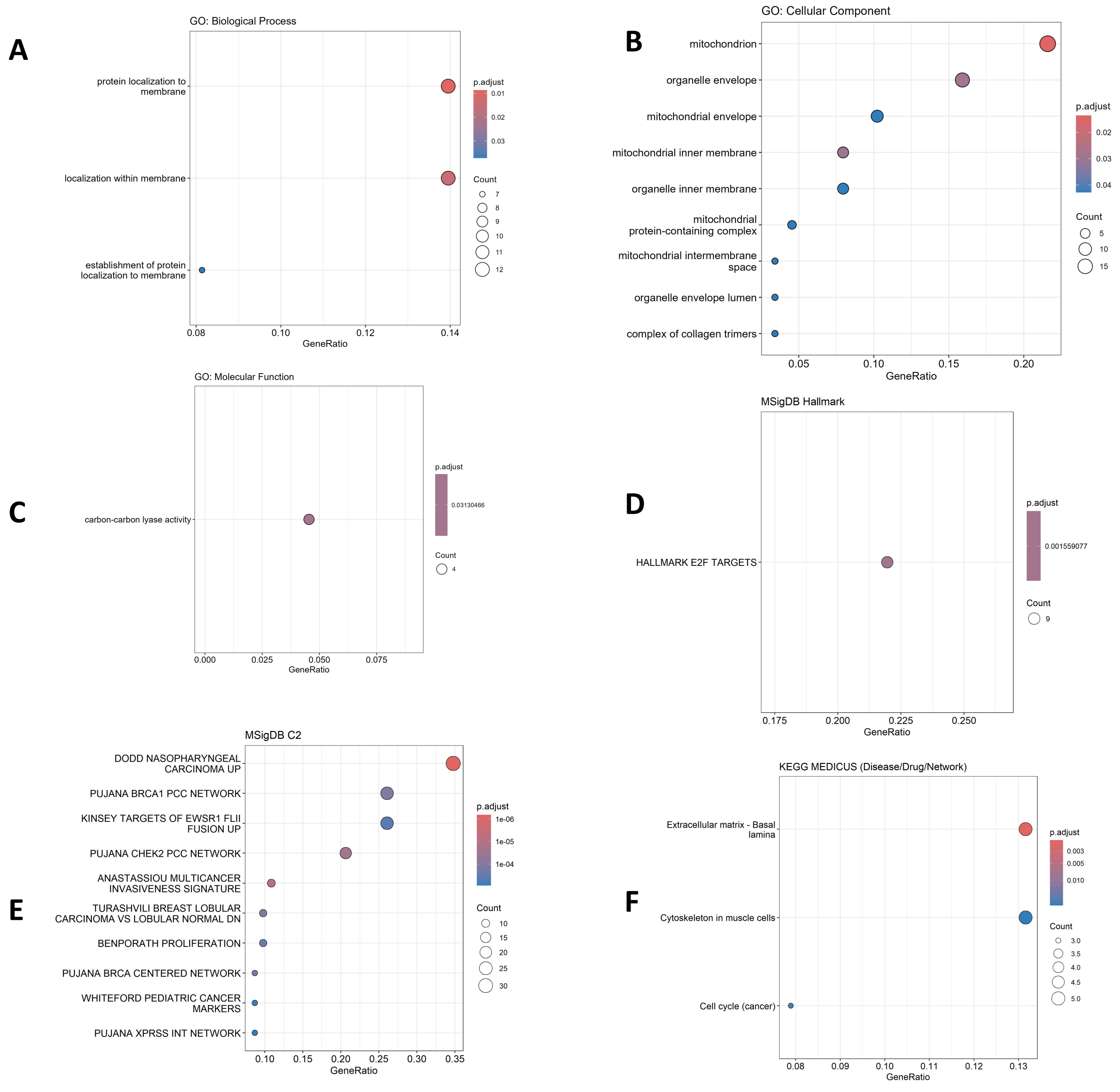


**Supplementary Figure 8**. Dotplots for over-representation analyses on MOFA extracted genes. ORA conducted on the 101 genes extracted from the MOFA model based on associations to treatment response. (a) Dotplot assessing enrichment for gene ontology biological processes. (b) Dotplot assessing enrichment for gene ontology cellular components. (c) Dotplot assessing enrichment for gene ontology molecular functions. (d) Dotplot assessing enrichment for MSigDB hallmarks. (e) Dotplot assessing enrichment for MSigDB C2 collections. (f) Dotplot assessing enrichment for KEGG medicus.

**A Combined Multi-Omics and Supervised Learning Strategy Uncovers an Epithelial Signature of Radiotherapy Response in Colorectal Cancer**

Reuben Kumar, Yujie He, Jiarui Zhou, S:CORT Consortium, Andrew D. Beggs, Deena M.A. Gendoo

#### Supplementary Figure 9

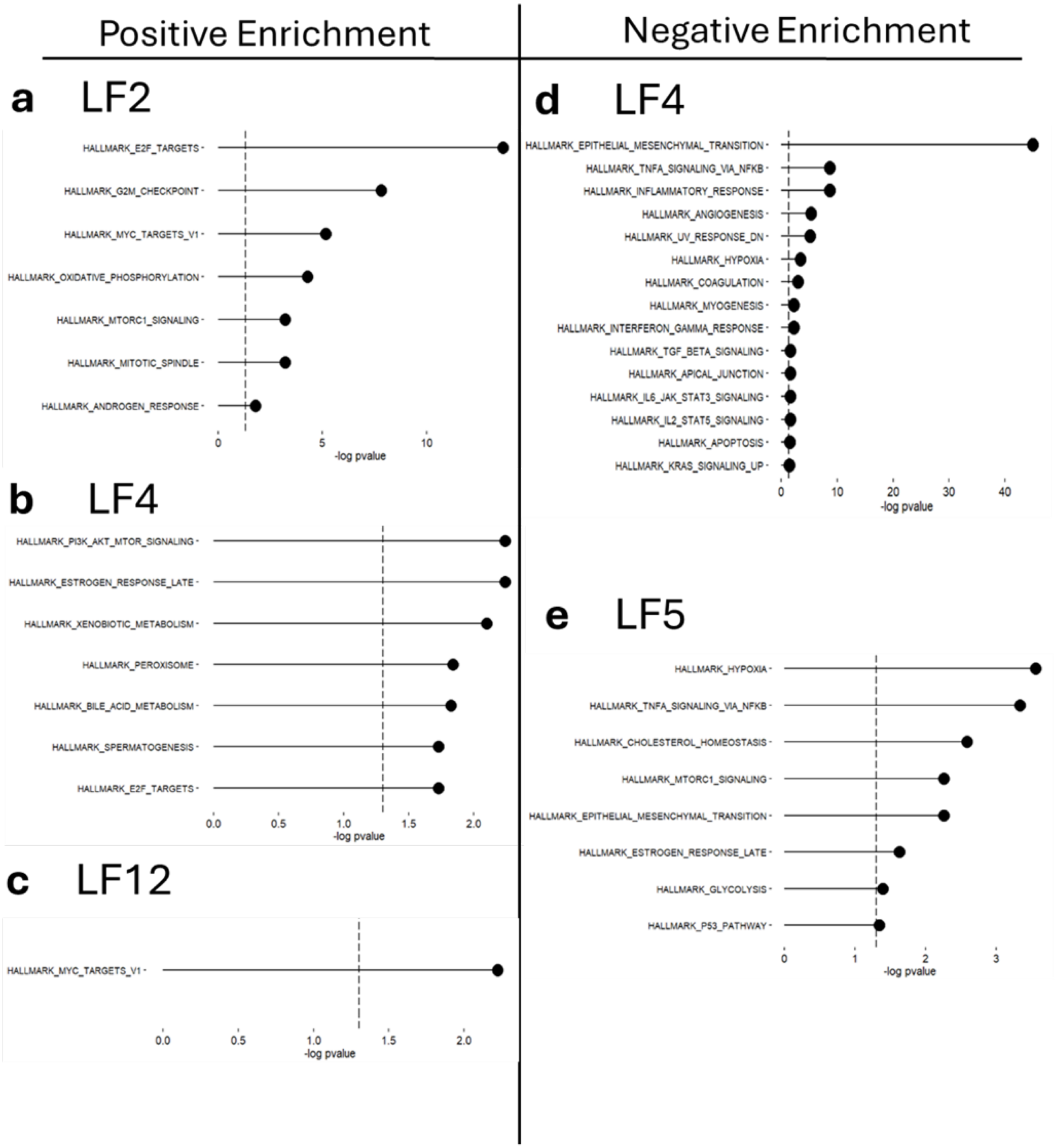


**Supplementary Figure 9.**
Gene set enrichment analysis (GSEA) of MOFA LFs correlated with treatment response. GSEA was performed separately on the positively and negatively loaded RNA features from the four LFs correlated to treatment response. Gene sets analysed were “Hallmark Gene Sets” from MSigDB. LFs 2,4 and 12 exhibited enrichment (a-c), whereas for the negatively loaded features only LFs 4 and 5 showed enrichment (d-e). The only significantly enriched gene set for LF12 was MYC targeting v1(c).

**A Combined Multi-Omics and Supervised Learning Strategy Uncovers an Epithelial Signature of Radiotherapy Response in Colorectal Cancer**

Reuben Kumar, Yujie He, Jiarui Zhou, S:CORT Consortium, Andrew D. Beggs, Deena M.A. Gendoo

#### Supplementary Figure 10

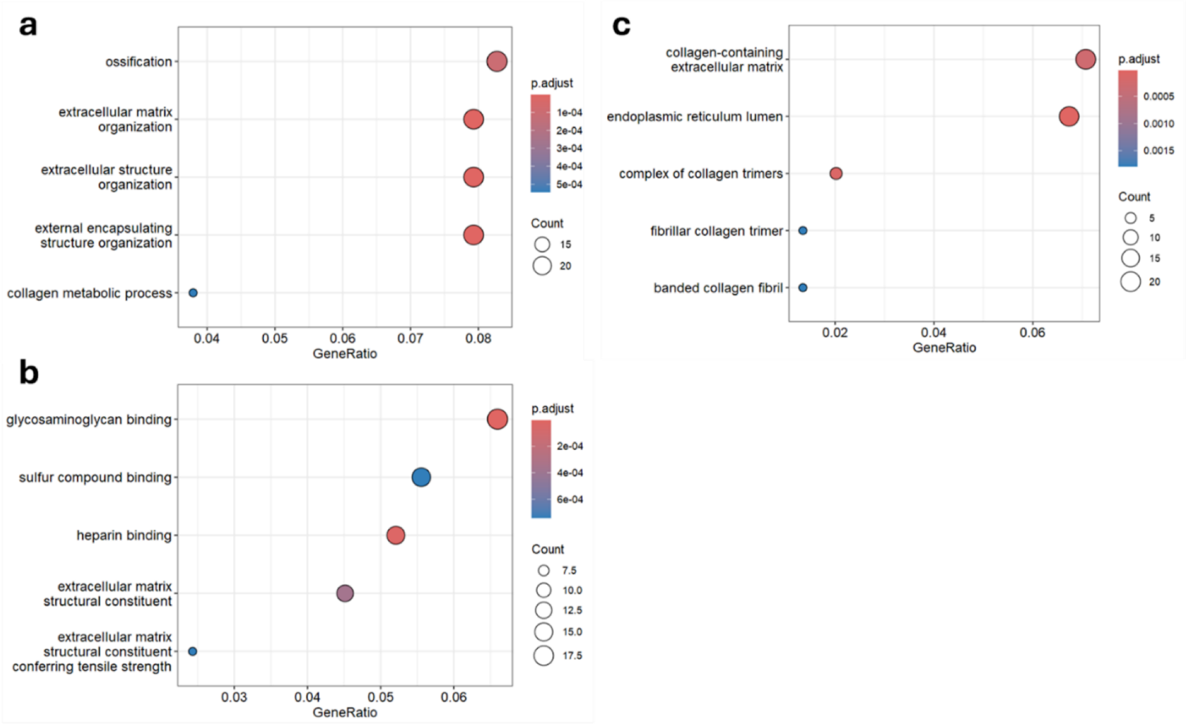


**Supplementary Figure 10.** Pathway enrichment analysis. Dot plots displaying significantly enriched pathways from the RNA features taken from MOFA LFs correlated to treatment response. Dot size indicates gene counts enriched in a pathway and colour represents adjusted p-values, with red showing higher significance. (a) Biological process ontology. (b) Molecular function ontology. (c) Cellular component ontology.
